# Adaptation to climate driven environments in a Patagonian suboscine passerine

**DOI:** 10.64898/2026.03.11.710818

**Authors:** Joseph Badji-Churchill, Martje Birker-Wegter, Maaike Versteegh, Rodrigo Vàsquez, Jan Komdeur

**Affiliations:** University of Groningen; Universidad de Chile

## Abstract

Climate change has altered global climatic conditions, which is affecting the reproductive strategies, offspring development, breeding biology and development of birds. We looked at the impact of different climatic variables (temperature, rainfall and wind speeds) before and during the nestling development phase on nestling development (i.e. nestling hatch weights, nestling growth rates and pre-fledging weights) in the Thorn-tailed Rayadito (Aphrastura spinicauda). We studied two populations. One is situated in a temperate rainforest on the northern border of Patagonia called Pucon which we studied in 2018 and 2019, with mild temperatures (12.5 degrees Celsius), high rainfall (636ml) and low wind speeds (6.3km/h). The other is in a sub-Antarctic old growth forest in southern Patagonia called Navarino island which we studied in 2018, 2019 and 2023, which is comparatively drier (138ml), colder (8.3 degrees Celsius) and has higher average wind speeds (16.6km/h). Embryonic development is key in ensuring individual future fitness. It is important that this is not interrupted and individuals are therefore vulnerable to damage during early development and it can have carry over effects into adulthood. Exposure to extreme climatic conditions can interrupt this development. Therefore, we expect to find that the climate during incubation to be important in predicting nestling hatch weights, growth rates and pre-fledgling weights. Climatic conditions are known to effect nestling development and extremes in climatic conditions have negative consequences on nestling development. We therefore expect that highly variable climatic conditions will have a negative effect on nestling development. We analysed populations separately because we expect populations to have developed different reaction norms to climatic factors. We found in both locations that hatching weights become lower each year, but growth rates and pre-fledging weights are unchanged. In Navarino, medium sized clutches produced the largest hatchlings whilst large and small clutches produced the smallest hatchlings and high or low rainfall during the egg laying and incubation phase produces smaller nestlings. No other climatic factors impacted hatch weights in Navarino. We also found that high or low average ambient temperatures during incubation and early nestling development in Navarino result in lower overall growth rates. Whilst in Pucon, lower rainfall and high or low wind speeds during incubation produce smaller hatchlings, but neither climatic nor biotics factors could explain growth rates in Pucon. We found pre-fledging weights could not be explained by climatic or biotics factors in either location. This is the first study of its kind to examine the environmental drivers of nestling hatch weights in birds in the wild. By better understanding how climate predicts nestling development, we can understand the potential future threats to fitness and development in birds with greater accuracy as conditions continue to change.

## INTRODUCTION

Climate change has severe impacts on the reproductive success of birds (Stevenson and Bryant, 2000; Carey, 2009). Variable climatic conditions interfere with annual mating and reproduction in many bird species. For example, inconsistent climatic conditions can cause birds to start breeding too early or too late and subsequently miss the seasonal food peak, resulting in less food being available for nestlings, leading to malnourishment (Visser, Both and Lambrechts, 2004; Bodey et al., 2021). Unfavourable climate can reduce the ability of adults to forage for food and sufficiently feed their young (Sicurella et al., 2015). Therefore, it is important to uncover what unfavourable climate looks like for different species. This question highlights important avenues of research in the behavioural plasticity of individuals. Behavioural plasticity is defined as an organisms’ ability to adapt to changing selection pressures like climate change (Tuero et al., 2018). For example, species time their breeding to coincide with peak food availability to give their offspring the best chance of survival. If the climate changes and the food peaks become earlier or later, adults need to adjust their breeding behaviour to the new food peak accordingly. There are gaps in our knowledge of how highly variable climate patterns under climate change affect behaviour and development. More specifically, literature on how climate can predict the hatching weights of nestlings in the field are, to the best of our knowledge, entirely absent. It is known that nestling growth rates from hatching to fledging are affected by climate, genetics, food abundance, sibling competition, habitat quality and parental investment (Gardner et al., 2011). But what is not known is when specifically, during the life cycle of a nestling (from embryo formation till fledging) does climate influence a nestlings growth rate. For example, is the climate during early or late development more important in predicting nestling growth? Or is the climate on the day nestlings hatch the most important in predicting growth rates throughout development? These questions are unanswered by current literature and is one of the main aims of this study. Early studies on the impact of climatic conditions over the breeding season of common swifts (*Apus apus)*, barn swallows (*Hirundo rustica*) and red-winged blackbirds (*Agelaius phoeniceus*) found that unfavourable climate are associated with low nestling growth rates and low pre-fledging weights (Lack, 1951 and Ricklefs, 1967). These findings are supported by more recent studies that have expanded into how nestlings respond to changing climate (Ruuskanen, Hsu and Nord, 2021; Garrido-Bautista et al., 2023). The review by Ruuskanen, Hsu and Nord, (2021) found that in many studies, there was no effect of climate on nestling growth rates, despite harsh and changing climatic conditions. They attribute this to plasticity in thermoregulatory behaviour. They stipulate that if individuals can adjust their thermoregulation, the effects of unfavourable climate may not be reflected in nestling development. Higher thermoregulatory plasticity to changing temperatures may mitigate potential impacts to development. However, what remains understudied is the delayed effect of climate on nestling growth rates. Insectivores time their breeding so that nestling hatching coincides with the peak insect abundance during the breeding season. Insect abundance is dictated by the climate as they rely on thermal triggers to initiate their breeding cycles (Müller et al., 2024). The studies in the review by Ruuskanen, Hsu and Nord, (2021) have not investigated the effects of climatic conditions before nestlings hatch which is when insect breeding cycles are triggered. This may explain the previous absence of an association between climatic factors and nestling growth rates.

The effects of climate on nestling development are species and habitat specific (Sauve, Friesen and Charmantier, 2021). An example from Mainwaring and Hartley, (2016) investigated ambient temperature, rainfall and daily average wind speeds on the nestling growth rates of blue tits (*Cyanistes caeruleus*) in the north of England. They measured nestling growth and observed that higher temperatures correlated with smaller nestlings. Another study on blue tits in the Mediterranean (Grosbois et al., 2006) found that temperatures which correlate with smaller nestlings in the north of England produce larger nestlings in the Mediterranean. These two examples demonstrate how the effects of climate on nestling development are habitat specific. The effect of climate on nestling growth can even differ within latitudes. One study by Bradbury et al., (2003) in England on common linnets (*Linaria cannabina*), chaffinches (*Fringilla coelebs*) and Eurasian skylarks (*Alauda arvensis*) found varying impacts of climate conditions on nestling growth. Chaffinches were unaffected by climate whereas skylark nestling growth increased with increasing minimum temperatures but declined with increasing maximum temperatures. Chaffinches feed exclusively on seeds and nuts, whilst skylarks are generalist feeders (Bradbury et al., 2003). The response of a species to unfavourable climate usually depends on their food source’s reaction to climate (Sauve, Friesen and Charmantier, 2021). Another study by Komdeur, (1996) on Seychelles warblers (*Acrocephalus sechellensis*) demonstrates this connection. In this study, rainfall predicted the food availability to warblers two months later, this triggered bird breeding after heavy rainfall as birds expected insect abundances to peak soon after, whereas on a separate island, food abundance was high year-round, so birds likewise bred all year round. Such a delayed effect has been found in other species. One example is from Facey et al., (2020) who found that more extreme temperature and rainfall in early nestling development were associated with poorer pre-fledging weights in barn swallows.

In this study, we investigate whether and how climate variations are associated with nestling hatch weights, growth rates and pre-fledging weights in the thorn-tailed Rayadito (*Aphrastura spinicauda*) over time. Previously, we have found strong associations of climate in the laying and incubation period to be highly relevant to pre-fledging haematocrit levels, where climate had a delayed effect on nestling haematocrit levels (Badji-Churchill et al., 2023). We explore if delayed climatic effects, i.e. ambient daily average temperatures, ambient daily minimum temperatures, ambient daily maximum temperatures, daily rainfall and daily average wind speeds can predict nestling hatch, growth and pre-fledgling weights. The thorn-tailed Rayadito is a common suboscine passerine across Chile and Argentina (Figure 1). It produces 1-2 clutches of 2-7 eggs each breeding season between September-January every year (Altamirano et al., 2015). It is useful to study nucleic species (locally common across large geographic distribution) such as the Rayadito because changes in behaviour, physiology or population dynamics can be indicative of a wider impact on other species, especially endemic or specialised ones, which is highly relevant to our study areas. By examining the influence of climate over time on nestling development, we hope to gain a better overall understanding of how birds respond to climate-based selection pressures. We use two populations of Rayaditos in two contrasting locations which vary in climatic conditions to investigate the association of climate variation and nestling development.

**Figure 1.**
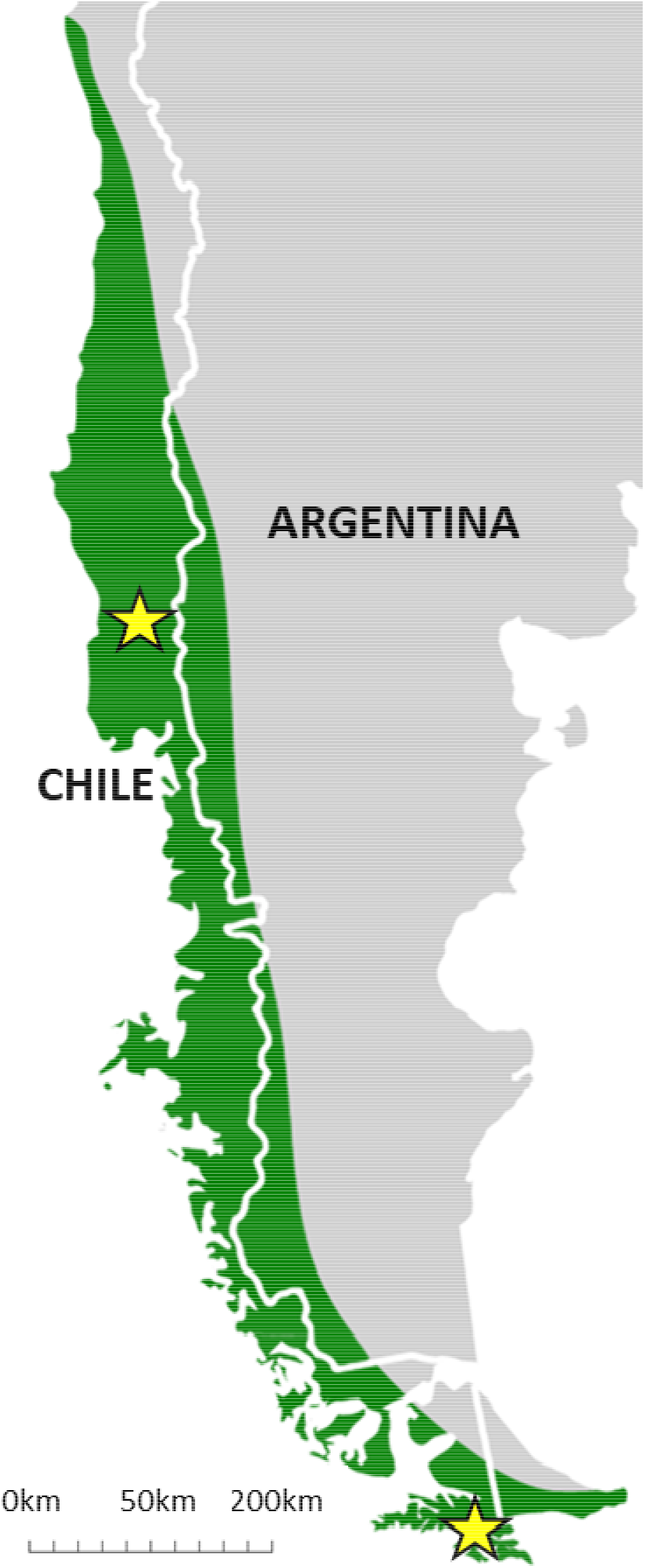
Locations of our two study sites. The yellow star in the north indicates the temperate rainforest (Pucón) and the yellow star in the south indicates the sub-Antarctic forest (Navarino Island). The green coverage shows the distribution of the thorn-tailed Rayadito. Map adapted from [https://birdsoftheworld.org].

The aim of this study is to explore what factors predict hatch weights, growth rates and pre-fledgling weights of thorn tailed Rayadito nestlings. We hypothesise that the climatic conditions at key developmental stages like egg laying, the initiation of full incubation and hatching etc. can predict the hatching weight, overall growth rate and pre-fledgling weights of nestlings during rearing. We predict weight and growth will be lower in more extreme conditions at these key stages (i.e. slower growth with very high or very low temperatures). There are three possible reasons for climate being able to predict hatch weights, growth rates and pre-fledging weights, firstly as discussed, climate mediates insect abundances and this has a direct effect on how much food nestlings receive. Secondly, more extreme climate dictates the behaviour of the parents, i.e. very wet or windy conditions may cause adults to forage less and so provide less food for nestlings. Finally, climate may have a direct effect on the nestlings themselves and cause thermoregulation stress.

## METHODS

### Study area and population

We studied two Rayadito populations in two different latitudes in Chile that breed in nest boxes (Figure 1). We collected data on Rayadito embryos during the breeding season (September to December) at two locations, Navarino island (54.932° S, 67.605° W) in 2018, 2019 and 2023 and Pucón, (39.272° S, 71.977° W) ca. 1800 km to the north in 2018 and 2019. The southern Navarino site was ca. 5km in radius, had around 200 nest boxes with 14.3% occupancy rate which were placed in 2002-3, in sub-Antarctic deciduous forest. The ambient average temperature over the breeding seasons in the years we collected data was 8.4°C (7.7°C - 9.2°C), total yearly rainfall averaged 138ml (87ml - 202ml) and wind speeds averaged 16.62km/h (15.95km/h - 17.80km/h) (INIA, 2025) (Table 1). Nest boxes in Navarino were dominated by deciduous species such as *Nothofagus antarctica, N. betuloides* and *N. pumilio*, whilst the understory was dominated by evergreen shrubs (*Berberis mycrophylla*). The Pucón study site had 240 nest boxes with 10.2% occupancy rate, spread out over ca. 10km radius which were placed in 2014. Pucón is dominated by a temperate rainforest and is comparatively wetter, warmer and less windy than Navarino. During the breeding season the temperature averaged 12.6°C (12.5°C - 12.6°C), there was an average total yearly rainfall of 636ml (512ml - 760ml) and the wind speeds averaged 6.3km/h (6.2km/h - 6.4km/h) (INIA, 2025) (Table 1). In Pucón, tree species were predominantly conifer-broadleaf species such as *Lophozonia obliqua, Nothofagus dombeyi, Laurelia sempervirens, Saxegothaea conspicua* and *Laureliopsis philippiana*. The understory composition was dominated by Bamboo (*Chusquea spp)*, evergreen shrubs (*Rhaphithamnus spinosus*) and flowering plants (*Azara spp*).

**Table 1.** The weather and food abundance (insect abundance) variables in Navarino and Pucón area (2018-2023). Data on Rayaditos was only collected in 2018, 2019 and 2023. NA = No data collected in those years. The average weather is followed by the (lowest/highest) average over the breeding season. Except for rainfall which is the total rainfall over the breeding season.

|  | NAVARINO |  |  |  |  |  | PUCON |  |
| --- | --- | --- | --- | --- | --- | --- | --- | --- |
|  | 2018 | 2019 | 2020 | 2021 | 2022 | 2023 | 2018 | 2019 |
| **Average Temperature (°C)** | 7.9<br>(4.9 / 10.7) | 8.0<br>(5.2 / 11) | 8.5<br>(6.0 / 10.9) | 8.9<br>(6.6 / 11.4) | 9.2<br>(6.9 / 11.5) | 7.7<br>(4.9 / 10.4) | 12.5<br>(11.8 / 13.2) | 12.6<br>(12 / 13.3) |
| **Minimum Temperature (°C)** | 3.1<br>(0.85 / 5.4) | 3.0<br>(0.72 / 5.0) | 3.5<br>(1.5 / 6) | 3.3<br>(0.90 / 5.2) | 3.6<br>(2.0 / 5.3) | 3.0<br>(2.4 / 3.6) | 5.8<br>(5.0 / 6.5) | 5.7<br>(5 / 6.5) |
| **Maximum Temperature (°C)** | 10.1<br>(7.8 / 12.8) | 10.1<br>(7.3 / 13.1) | 10.6<br>(8.7 / 13.0) | 11.4<br>(9.1 / 13.9) | 11.6<br>(9.5 / 13.6) | 9.8<br>(7.1 / 12.1) | 18.8<br>(17.7 / 19.7) | 19<br>(18.1 / 19.9) |
| **Total Rainfall (ml)** | 162.8 | 202.3 | 153.5 | 117 | 87 | 104 | 760 | 512 |
| **Average Wind Speed (km/h)** | 17.8<br>(16.2 / 19.4) | 17<br>(15.6 / 18.5) | 15.95<br>(14.26 / 17.64) | 16.39<br>(12.03 / 20.75) | 16.48<br>(14.04 / 18.92) | 16.1<br>(14.7 / 17.5) | 6.4<br>(6.0 / 6.8) | 6.2<br>(5.9 / 6.6) |
| **Insect abundance**<br>(average Insect numbers over breeding season) | 27.93 | 27.35 | NA | NA | NA | 12.41 | 7.2 | 5.19 |

### Data collection

Data were collected in three years in Navarino and Pucón from September 2018 – January 2019, September 2019 – January 2020 and in Navarino only in October 2023 – December 2023, during the Rayadito breeding season. Rayaditos in Pucón start the breeding season in mid-September until mid-January. In Navarino, the breeding season starts mid-October until the end of January. Data will be referred to from here as the 2018, 2019 and 2023 years, respectively. We checked all nest boxes every 4 days to track the nest building phase and ensure we did not miss the first egg laying of each occupied nest box. Subsequently, if activity was observed, we visited the nest box every 2 days.

We sampled 343 nestlings from 104 nests over three years in two locations in Chile. We collected data from 96 nestlings in 2018, with 45 nestlings from 17 nests in Navarino and 51 nestlings from 25 nests in Pucón, respectively. We measured 173 nestlings in 2019, with 139 in Navarino from 32 nests and 34 nestlings in Pucón from 13 nests, respectively. Finally, in 2023 we collected data from 74 nestlings in Navarino from 17 nests.

### Nestling morphometrics

The Rayadito is a synchronously breeding species and all eggs hatch within a 24-hour period (Moreno et al., 2005). When the clutch hatched, this was recorded as day 1 (D1), which was standardised as the day of hatching. On D1, we measured the body mass (g) to the nearest 0.01g, using digital scales. To differentiate between individual chicks, we clipped the tip of a toenail of each individual to tell them apart. We then returned to weigh them on Day 4 (D4) and on day 8 (D8), when in addition we ringed each nestling. We weighed them again on day 12 (D12) and day 16 (D16). We then checked nests on day 24 (D24) to ensure all nestlings had fledged successfully as nestlings fledge between D18 - D22 (Moreno et al., 2005; Espíndola-Hernández et al., 2017).

### Climatic and insect measurements

Temperatures (degrees Celsius), rainfall (ml) and wind speed (km/h) data were taken from meteorological stations in both sites (Table 1). In Navarino, the weather station was a maximum of 5 km away from the nest boxes and in Pucón the weather station was located a maximum of 7 km from the nest boxes (INIA, 2025). We collected daily ambient average temperature, daily ambient minimum temperature, daily ambient maximum temperature, total daily rainfall and daily average wind speed, from 1^st^ September each year up until the last nestling measurement was taken. The daily ambient average temperature and daily average wind speed was calculated with hourly temperatures from 6 am - 6 pm, but we the full 24 hours to calculate total rainfall and the minimum/maximum temperature.

Insect counts began when the first egg was laid at each location and were then repeated once every 6 days in all years. In 2018 in Pucón, insect counts began 23/09/2018 and finished on 22/01/2019, and in 2019 they began 25/09/2019 and finished on 03/01/2020. Insect counts on Navarino in 2018 began 25/09/2018 and finished on 10/01/2019, in 2019 they began 14/10/2019 and finished on 09/01/2020 and then in 2023 they began 04/10/2023 and finished on 13/12/2023. In all years, we used four random trees of random species in each location to sample insect abundance every 6 days. The random tree was selected by randomly selecting a spot on the GPS device (accurate to 3 metres) in the same area where the nest boxes were and used the tree closest to the random GPS spot. The insect counts were conducted by using a 3-sided cloth sheet (80 cm x 80 cm x 80 cm) attached to 3 poles, which was held up underneath a branch that was struck 10 times with a stick to dislodge anything clinging to the branches. After striking the branch, the sheet was lowered, and the insects and spiders were counted for total abundance, giving a weekly average from all trees in different areas (see Leather, 2008).

### Statistical analyses

All statistics were carried out using R in RStudio v3.5.3 statistical software (Posit team, 2026). We conducted a mix of analyses using climwin models and LMERs (Linear Mixed Effect Models).

We wanted to know the specific days in which ambient average, minimum and maximum temperature, rainfall and wind speeds correlated with nestling hatch, growth and pre-fledging weight. This will tell us if climate has a delayed effect on nestling weight and growth. We did this using the ‘climwin’ package. This package correlates different climate windows on a response variable and evaluates best fit based on Akaike Information Criterion (AICc) (see Bailey and Van de Pol, 2016). We used hatching weight (g) (D1), pre-fledging weight (g) (D16) and growth per day (g) as our response variables, with daily average temperature, daily minimum temperature, daily maximum temperature, daily total rainfall and daily average wind speed as our explanatory variables with year as our baseline variable and nest box as a random variable (Bailey and Van de Pol, 2016).

The ‘slidingwin’ function in the climwin package was used on our response variables with the average, minimum and maximum temperatures (°C), total rainfall per day (ml) and daily average wind speed (km/h) in two separate models (linear and quadratic) to identify the time-window where climatic factors explained weight and growth best. This package calculates several models considering multiple windows relating a response variable to the climatic factors at different time resolutions. Then, the best model is selected through an information-theoretic approach and randomization tests are computed to establish the significance of the selected model. We used climatic data from 20-30 days prior to measurements, which covers the incubation period (~14 days), egg laying (~8 days), hatching and brooding periods (~16 days). We ran 1000 random models to determine that our best window is significantly different from random. If a reliably significant relationship was found, this data was then extracted for further LMERs with other predictor variables. We analysed both locations separately in this analysis because we expect populations to have developed different reaction norms to climatic factors.

In our LMERs we used several predictors on the three response variables (hatching weight, growth rate and pre-fledging weight. In Navarino, for hatching weights, we used the climate variables daily rainfall (7-18 days before hatching), and for growth rates we used the daily average temperature (13-26 days before fledging). In Pucón, for hatching weights we used daily minimum temperatures (17.5-19 days before hatching), daily total rainfall (10-15 days before hatching) and daily average wind speeds (10-11 days before hatching).

For pre-fledgling weight in Navarino, growth rate and pre-fledgling weight in Pucón, climwin identified no climate variables as important so we did not include any in our LMERs. Other predictor variables we used were insect abundance, incubation length, and clutch size or brood size, hatching weight and growth rates. We used year as covariate and nest box as a random factor in all models. We did not include “hatching date” as a covariate in our analyses because all the data we recorded are the reasons why growth rates may diminish towards the end of the breeding season (see Badji-Churchill et al., 2023).

## RESULTS

### Delayed climatic impacts on nestling hatching weight, fledging weight and growth rates

Mean average hatching weight in Navarino was 1.90g, sd = 0.29, (306 nestlings in 2018, 2019 and 2023). Mean average hatching weights in Pucón were 1.77g, sd = 0.35 (171 nestlings in 2018 and 2019). Nestlings were significantly smaller at hatching in Pucón compared to Navarino (*Est* = −0.13, *sd* = 0.03, *P* = <0.01). The average incubation length in Navarino was 16.6 days (rounded up to 17), sd = 1.30, and in Pucón it was 16.3 days (rounded down to 16), sd = 1.14. In Navarino, climwin identified daily total rainfall 7-18 days before hatching as the only predictor for hatching weights in a quadratic effect (Figure 2). More specifically, when rainfall was light or heavy the day before full-time incubation to 10 days of full-time incubation (11-day window) hatch weights were smaller. The climwin results in Pucón showed that daily rainfall 10-15 days before hatching, the daily average wind speeds 10-11 days before hatching and the daily minimum temperatures 17.5-19 days before hatching predicted hatching weights (Figures 2 and 3). However, in the case of daily minimum temperatures, further analysis reveal that this had no impact on nestling hatch weights (Table 2).

**Table 2.**
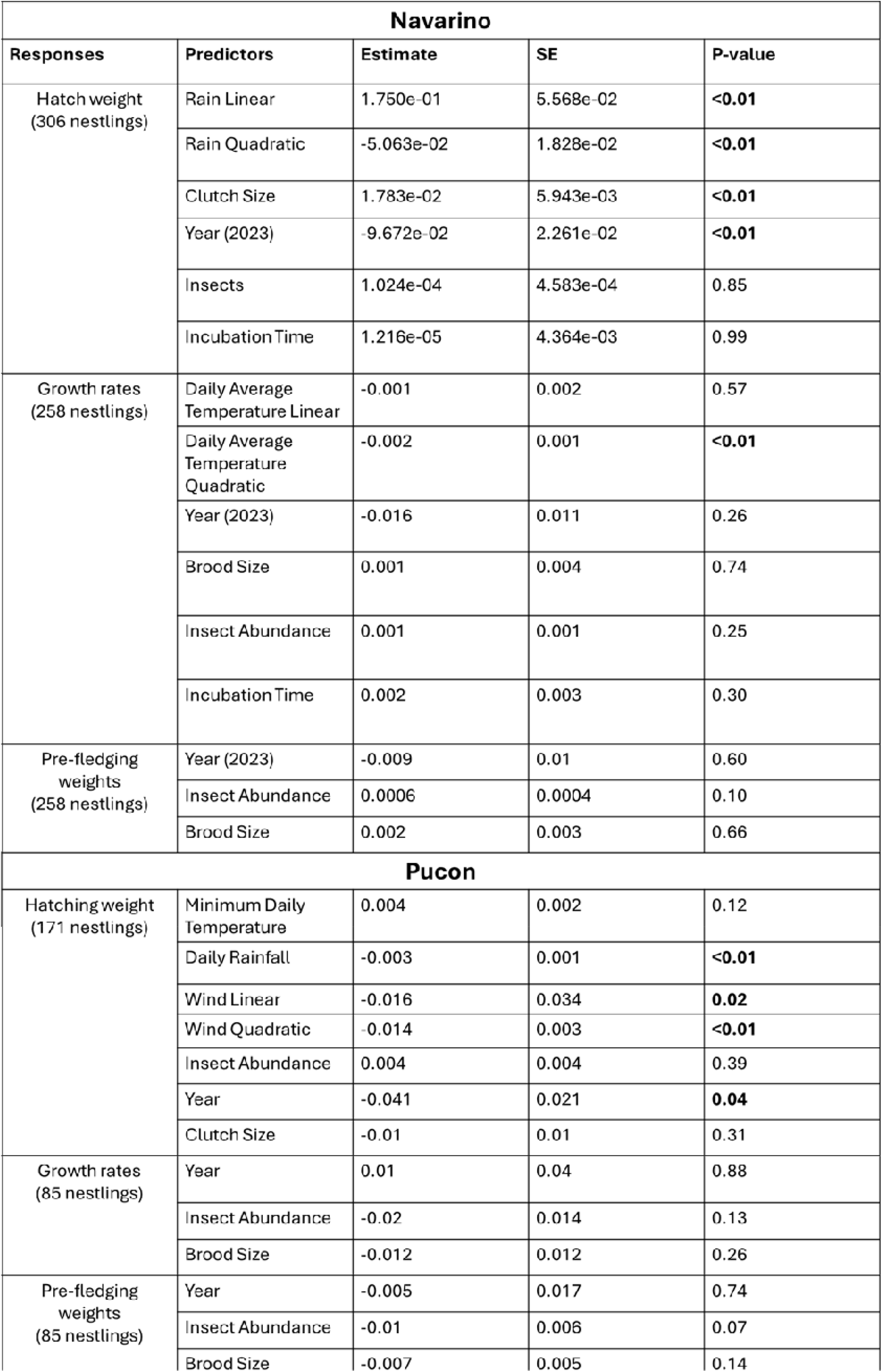
The LMERs of the hatching weights, growth rates and pre-fledgling weights of thorn-tailed Rayadito in Pucón and Navarino.

**Figure 2.**
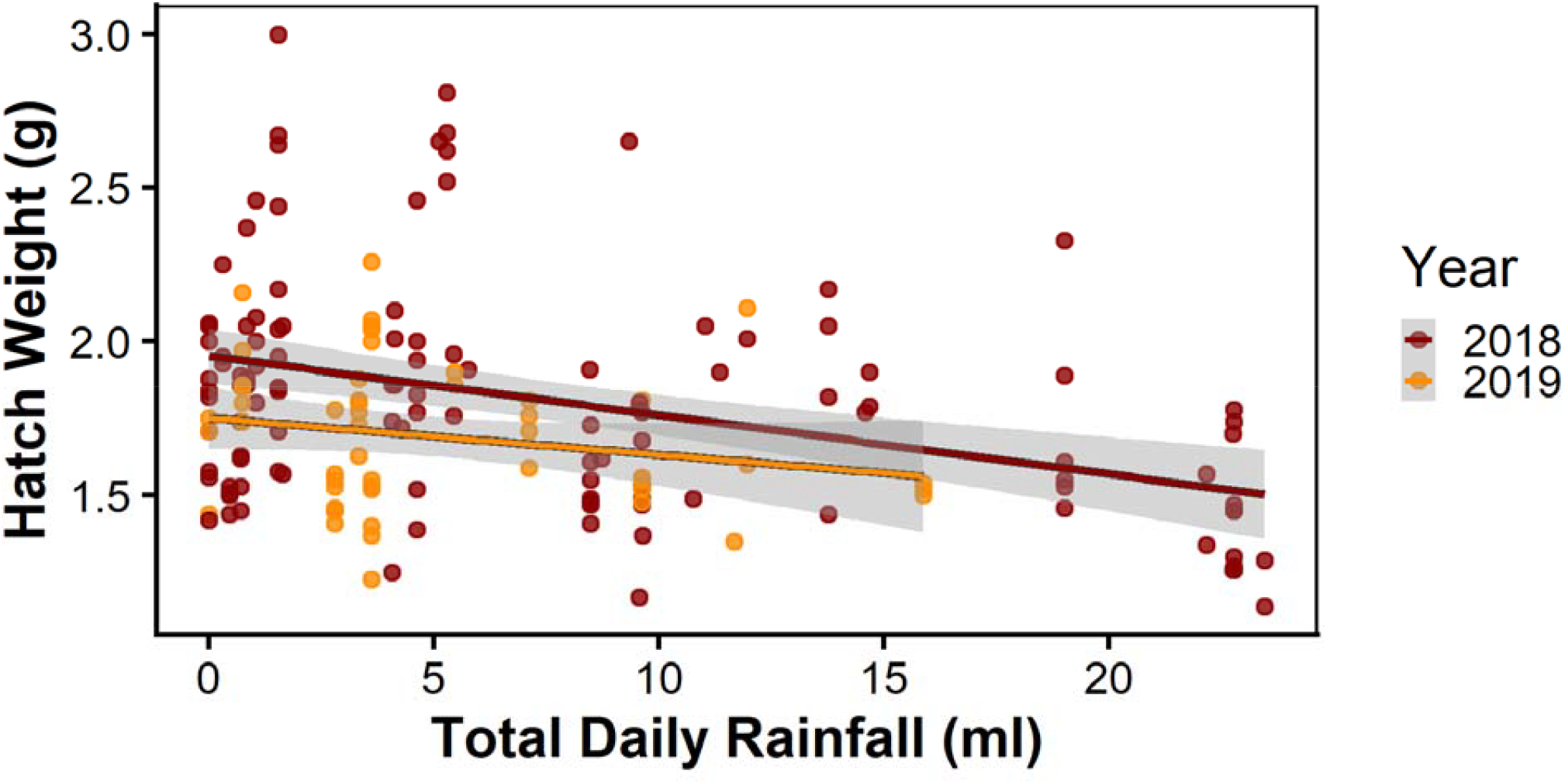
Relationship between daily total rainfall during early incubation (10-15 days before hatching) and thorn-tailed Rayadito nestling hatch weights in Pucón (2018, N = 120. 2019, N = 51). For statistical analyses see (Table 2).

In the LMERs, rainfall had a negative linear effect on nestling hatch weights where more rainfall resulted in smaller nestlings (Figure 2). More specifically, when rainfall was heavy from the first day of full incubation to the 6^th^ day of full incubation (5-day window), hatch weights were smaller. The daily average wind speed had a quadratic effect on nestling hatch weights where low or high wind speeds resulted in smaller hatchlings and moderate wind speeds resulted in the largest hatchlings (Figure 3). More specifically, when wind speeds were high or low from the 5^th^ – 6^th^ day of full incubation (2-day window), hatch weights were smaller.

**Figure 3.**
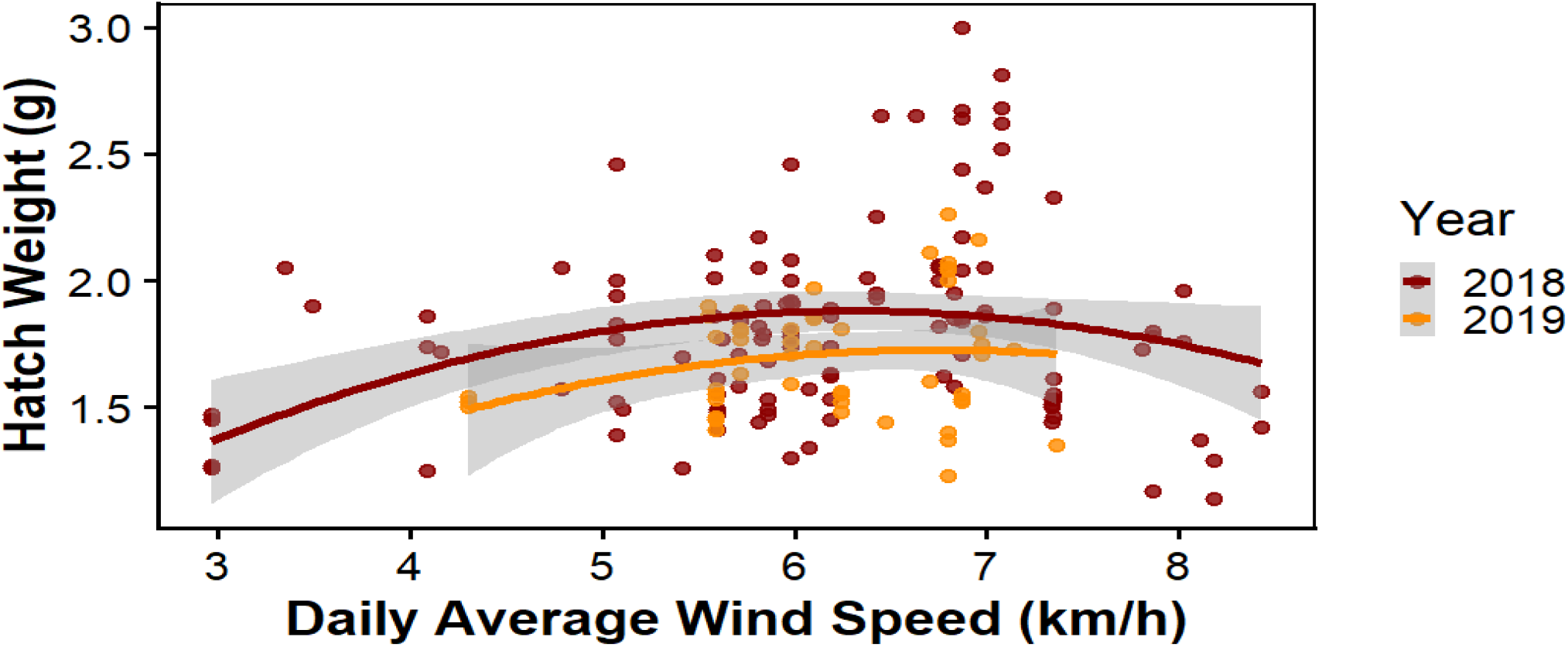
Relationship between daily average wind speed during early incubation (10-11 days before hatching) and thorn-tailed Rayadito nestling hatch weight in Pucón (2018, N = 120. 2019, N = 51). For statistical analyses see (Table 2).

Mean average nestling growth per day in Navarino was 0.91g/day, sd = 0.09, (258 nestlings in 2018, 2019 and 2023). Mean average nestling growth per day in Pucón was 0.82g, sd = 0.10 (85 nestlings in 2018 and 2019). Pucón had significantly lower nestling growth per day (*Est* = −0.09, *sd* = 0.005, *P* = <0.01). In contrast, in Navarino, the daily average temperature 13-26 days before the last day of growth measurements predicted overall growth rates of nestlings in a quadratic relationship. More specifically, when ambient average temperatures were high or low from the 6^th^ day of full incubation to when nestlings are 3 days old (13-day window) growth rates were lower (Figure 6). Average pre-fledging weight in Navarino was 15.5g, sd = 1.4 (258 nestlings in 2018, 2019 and 2023). In Pucón, no climatic variables could predict the growth rates of nestlings. There were no correlations between climate and prefledging weights in Navarino or Pucón. Average pre-fledging weight in Pucón was 14.1g, sd = 1.4 (85 nestlings in Pucón in 2018 and 2019). Pucón had significantly smaller pre-fledging weights (Est = - 1.45, sd = 0.08, P = <0.01).

### Other impacts on hatching and fledging weight and growth rates

In Navarino, clutch size influenced hatch weights (Table 2 and Figure 5). Small and large clutches produce smaller nestlings whilst clutches of three produce the largest nestlings. Insect abundance and incubation time had no effect on nestling hatch weights. In Pucón, neither clutch size nor insect abundance influenced hatching weights. Similarly, in Navarino and Pucón, insect abundance and brood size had no impact on nestling growth rates or pre-fledgling weights. In Navarino, we observed that hatch weights of Rayadito nestlings declined over the years, with the smallest on record in 2023, but growth rates and pre-fledgling weights remained unchanged (Table 2). In some cases, the climate differed between years in Navarino.

## DISCUSSION

In this study we investigated the effect of climatic factors (daily average ambient temperature, daily maximum temperature, daily minimum temperature, daily total rainfall and daily average wind speeds) on the hatching weights, growth rates and pre-fledging weights of the non-migratory thorn tailed Rayadito at two locations in Chile, which varied markedly with respect to climatic and habitat conditions. We found that climate had a delayed effect on nestling development in Rayaditos, often with climatic factors during incubation and early development predicting nestling hatch weights and growth rates. Nestlings in Navarino were 7.2% larger at hatching, had 9.6% higher daily growth rates and were 9.2% larger at pre-fledging compared to Pucón.

Non-climatic factors had a limited impact on nestling development with only clutch size in Navarino being able to predict hatch weights; small and large clutches had smaller hatchlings whilst clutches of three produced the largest nestlings. Otherwise, clutch/brood size, food abundance and incubation length had no effect on nestling development. Rainfall had a delayed negative linear effect on hatch weights in Pucón, where heavy rainfall from the first day of full incubation to the 6^th^ day of full incubation (5-day window) produced smaller hatchlings (Figure 2). Daily average wind speeds had a delayed quadratic effect on hatch weights in Pucón, where high or low wind speeds from the 5^th^ – 6^th^ day of full incubation (2-day window) resulted in smaller hatchlings (Figure 3). Daily total rainfall had a delayed quadratic effect on hatching weight in Navarino, where light or heavy rainfall the day before full-time incubation to 10 days of full-time incubation (11-day window) resulted in smaller hatchlings (Figure 4). Growth rates and pre-fledging weights were complicated. Daily average ambient temperature had a delayed quadratic effect on growth weights in Navarino where high or low ambient average temperatures from the 6^th^ day of full incubation to when nestlings are 3 days old (13-day window) growth rates were lower (Figure 6). Although climatic factors do have an impact on nestling development in some cases, in many cases it has no effect. For example, no climatic factor had an influence on pre-fledging weights in either location. The most likely reason for this is that parents make a trade-off during nestling development by investing more into parental care through extra feeding to ensure pre-fledging weights are optimised. This is likely why we find that climatic factors have the most influence on hatch weights (early in development) and then further into development, the influence of climate fades as these effects are mitigated by increased parental investment. The question following this is what the threshold of increased parental investment is, as parental investment cannot increase exponentially to mitigate the consequences of climate change.

**Figure 4.**
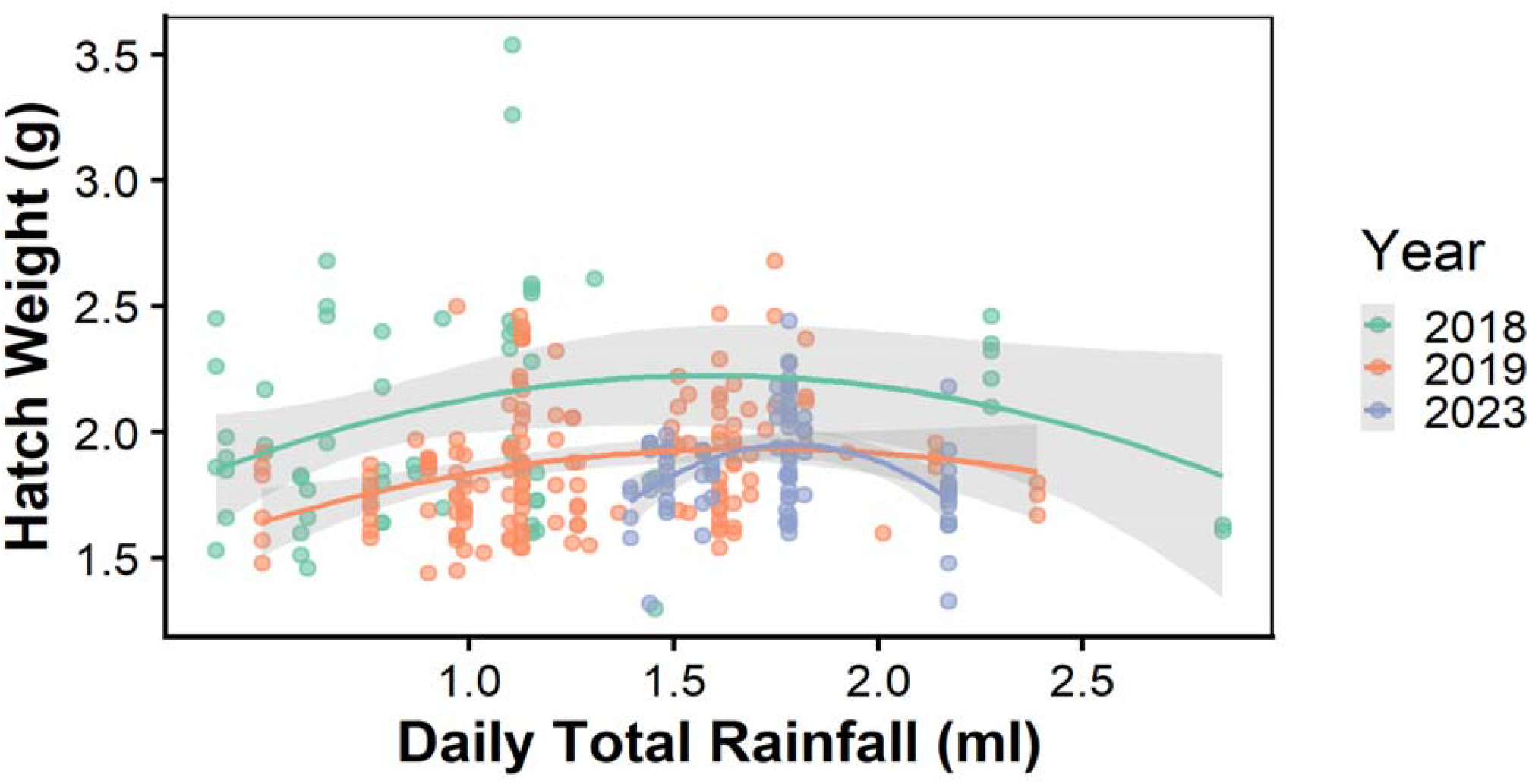
Relationship between the total daily rainfall (ml) during incubation/egg laying period (7-21 days before hatching) and thorn-tailed Rayadito nestling hatch weights in Navarino (2018, N = 66, 2019, N = 156, 2023, N = 84).

**Figure 5.**
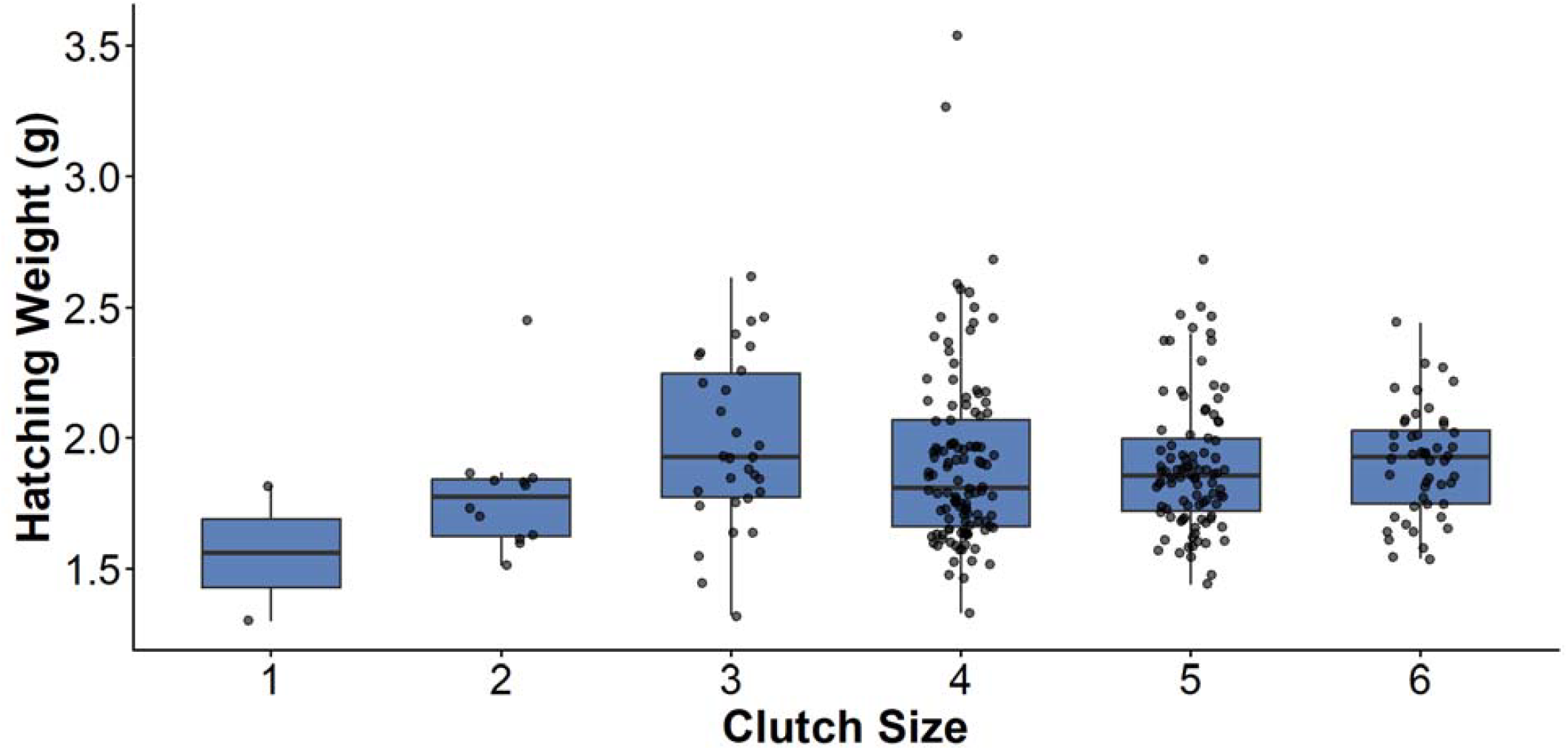
Relationship between hatch weight and clutch size of Rayaditos in Navarino (2018, N = 66, 2019, N = 156, 2023, N = 84).

### Location based differences in responses to climatic factors

Rayaditos have adapted differently to the different climates and specific habitats they exist in. For example, wind speeds are comparatively high in Navarino and relatively low in Pucón. Birds have evolved being exposed to consistently high wind speeds in Navarino and as such we may not find an effect of high wind speeds on nestling weight or growth. However, in Pucón, where wind speeds are lower and more stable, birds may not have adapted to dealing with dramatic changes in wind speeds. We found that wind speeds in Pucón have a significant quadratic effect on nestling hatch weights; with high and low wind speeds corelating with lower hatch weights and moderate wind speeds correlating with the largest hatch weights. When these locations are combined, we find a strong negative linear relationship between wind speeds and hatch weights, which changes the way we perceive the effect of wind speeds on hatch weights. This is not an accurate reflection of the complexity of the relationship between climate and development in this species across its different habitats. Analysing locations separately represents the observed patterns more accurately.

### Delayed climatic impacts on hatching and fledging weight and growth rates

#### Hatching weights

The effects of climatic variables on nestling hatch weights in our study are many and varied. Climate had a clear effect on hatch weights in both locations. The total daily rainfall and daily average wind speeds during incubation were found to predict hatch weights in Pucón. Whilst daily ambient average temperatures, daily maximum and minimum could not predict nestling hatch weights.

To our knowledge there are no other published studies that explore the delayed effect of climatic variables on nestling hatching weights. The only studies are those investigating the effects of nest temperature during incubation through observations and experiments on hatching weights (Heenan, 2013; Vaugoyeau, Meylan and Biard, 2017). Internal nest temperatures and ambient temperatures are often independent (Coe et al., 2015). One study from Vaugoyeau, Meylan and Biard, (2017) experimentally heated nests with heat pads, artificially increasing the incubation temperature, and they found that heating the nest during incubation caused higher nestling weight at hatching, but nestling weight gain was slower, which resulted in similar fledgling weights with control groups. However, the key difference is that this study focuses on internal nest temperatures which are different from external atmospheric temperatures. What makes our study unique is that we used the ambient atmospheric temperatures as our predictor variable instead of nest temperature.

In Navarino, we found that low and high rainfall during the egg laying and incubation period resulted in smaller hatchlings. Rain had a quadratic effect on hatch weights where low and high rainfall correlated with smaller hatch weights, and moderate rainfall correlated with larger hatch weights. To our knowledge there are no studies reporting on the impact of rainfall on nestling hatch weights and only a few studies look at rainfall specifically as a predictor of nestling growth, but there is evidence that moderate rainfall encourages greater nestling growth. A study from Busch et al., (2011) found a quadratic relationship between rainfall and nestling growth in song wrens (*Cyphorhinus phaeocephalus*) where moderate rainfall was associated with the higher nestling growth. The reason for this was that moderate rainfall stimulates caterpillar movement (the birds main food source) and therefore increases food availability for foraging parents which leads to more food for nestlings resulting in higher nestling growth. However, this would not explain why moderate rainfall during incubation would lead to larger hatchlings in our study. This is a novel finding, and we hypothesize that rainfall influenced the incubation patterns of adults who spent more time in the nest incubating when rainfall was high which resulted in faster development and smaller hatchlings. This hypothesis is backed up by a study from Coe et al., (2015) who found that female tree swallows (*Tachycineta bicolor*) incubate eggs more often and for longer time periods in rainy conditions, although, in this study there was an interactive effect with ambient temperature, where warmer and wetter conditions led to less incubation and cold and wet conditions led to more incubation. In both our sites, higher rainfall linearly correlates with lower temperatures, so this explains how and why low or high rainfall during the incubation period would influence hatching weights. There is evidence that faster embryonic development (caused by more incubating) results in smaller hatchlings (Vedder et al., 2018; Stier, Metcalfe and Monaghan, 2020). However, this does not fully explain why hatchlings were smaller when rainfall was low during the incubation period. We further hypothesize that in Navarino, where lower rainfall correlated with higher ambient average temperatures, this increased internal incubation temperatures and accelerated embryonic growth, which resulted in smaller hatchlings. These results need to be followed up with further observations from other areas and with experimental manipulations of ambient temperature and rainfall to establish if there is a common pattern or if this finding at the Navarino site is unique. There was a difference between the effect climate had on hatching weights between Navarino and Pucón. Daily total rainfall influenced hatch weights in Navarino and Pucón, but it had a linear effect in Pucón and a quadratic effect in Navarino. This difference can be explained by the difference in rainfall between the two locations, as Navarino is very dry in comparison to Pucón (Table 1). The difference can be explained by the same reason that wind speed had a different effect (see “Location based differences in responses to climatic factors”, above). Rayaditos in Navarino are adapted to dry conditions, so small increases in rainfall may have a disproportionate effect on nestling development, which would explain the quadratic effect we found. Whilst Rayaditos in Pucón have adapted to comparatively heavy rainfall. Rayaditos in Pucón prefer rainfall that would be considered heavy in Navarino, so only very high extremes in rainfall can begin to have a negative effect on Rayadito development, which is likely why we find a linear effect. Additionally, as mentioned, wind influenced hatch weights in Pucón but not in Navarino (see “Location based differences in responses to climatic factors”, above for explanation).

#### Growth rates

The effects of climate variables on nestling growth rates are complex and nuanced (Facey et al., 2020; Sauve, Friesen, and Charmantier, 2021: Sauve et al., 2022). For example, Facey et al., (2020) found in barn swallows that higher ambient average temperature during early nestling phases predicted lower pre-fledging body weight, but the strength of this effect was mediated by rainfall and wind speeds. When it was drier and windier, nestlings were heavier, whilst nestlings were lighter in wetter and calmer conditions. In addition, Sauve et al., (2022) found the opposite in Kittiwakes, where warmer air temperatures predicted heavier and faster growing nestlings. There is variation in responses to climatic factors between species as seen here because different species have evolved differently to the same selection pressures. In Pucón, no climatic variables predicted nestling growth rates. On the other hand, in Navarino, when ambient average temperatures were high or low from the 6^th^ day of full incubation to when nestlings are 3 days old (13-day window) growth rates were lower (Figure 6). The effects of climate on growth rates of nestlings have been well studied, and our findings correspond with other studies (Gardner et al., 2011; Mainwaring and Hartley, 2016; Sauve, Friesen, and Charmantier, 2021; Riggio et al., 2023). However, only one study carried out a similar analysis (using climwin) on the delayed effects of temperature on nestling growth rates on alpine swifts (*Tachymarptis melba*) (Masoero et al., 2024). Ambient average temperature has a similar quadratic effect on nestling growth rates in alpine swifts as in our study. However, in this case the ambient temperature 2 days before the body weight measurement was taken was identified by climwin as the most important window. Whilst the ambient temperature 13-26 days before the body weight measurement was taken was identified by climwin as the most important window in our study. However, this study did not look at climate before hatching which may explain the difference between these two findings. It is encouraging that this study found the same relationship with ambient temperature through the same statistical approach and will help us build theoretical models on the response of population dynamics to climate change in the future.

**Figure 6.**
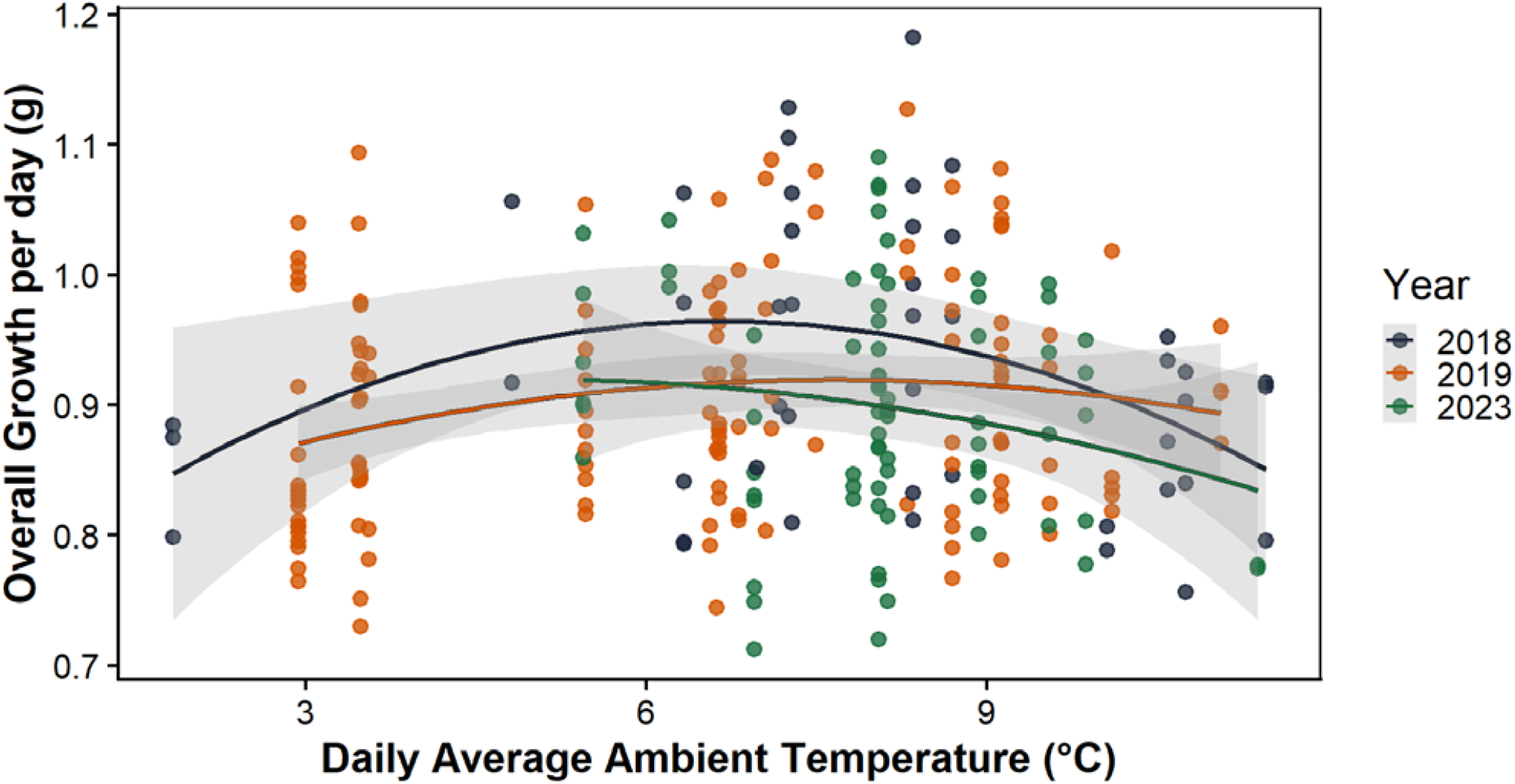
Relationship between daily average ambient temperatures during incubation period and early development (up to when nestlings are 3 days old) and thorn tailed Rayadito nestling growth rates in Navarino (2018, N = 45, 2019, N = 139, 2023, N = 74).

#### Nestling pre-fledgling weights

We found no effects of ambient temperature, rainfall or wind speeds on the pre-fledgling weights of nestlings, nor could biotic factors explain this, we even find that hatching weights did not predict pre-fledging weights as you may expect larger hatchlings to be larger at pre-fledging. The reasons for this are because parents increase their parental investment by feeding more when nestlings hatch smaller, which is clear mitigating the effect. This is further evidenced as nestling growth rates predict pre-fledging weights (Est = 0.41, SE = 0.01, P = <0.01), where nestlings who grew more were larger at pre-fledging.

#### Long-term development implications of climatic impacts on development

Research shows that fast development and shorter incubation periods lead to smaller hatchlings but longer telomere lengths and the potential for longer life spans. One study by Vedder et al., (2018) incubated common tern (*Sterna hirundo*) eggs under different incubation temperatures and extracted blood samples at hatching to measure telomere base pair lengths. Telomeres are non-coding DNA regions at the tip of chromosomes. Longer telomere lengths are associated with longer life spans in birds and this study found that higher incubation temperatures caused shorter incubation periods (and faster development) which resulted in longer telomere lengths in individuals, but who were smaller at hatching. This is supported by other studies in Japanese quails (*Coturnix japonica*) (Stier, Metcalfe and Monaghan, 2020) and purple-crowned fairy-wrens (*malurus coronatus*) (Eastwood et al., 2022). Based on these findings, if wetter, colder and windier conditions lead to more incubating by adults, this could lead to faster embryonic development and shorter incubation periods. This can result in smaller hatchlings, which may be disadvantageous in the short term, especially for nestlings in Navarino, where being large is essential to surviving the first winter. However, this may be advantageous in the long-term if smaller nestlings have longer telomere lengths and potentially longer life spans.

#### Biotic impacts on hatching and fledging weight and growth rates

Climatic factors influence hatching weights and growth rates in Rayaditos, but biotic factors such as clutch size, brood size, food abundance and habitat quality can equally have an impact on nestling development. Clutch and brood size can be important contributing factors to nestling development (Nur, 1984; Sicurella et al., 2015; Čech and Čech, 2017). In Navarino clutch sizes are larger than in Pucón and range from 1-6 eggs, with a mean clutch size of 4.5 eggs (compared to 3.4 eggs in Pucón) and has a significant quadratic relationship with hatching weight in Navarino. Small and large clutches in Navarino produce smaller hatchlings and clutches of three produce the largest hatchlings. Studies on European starlings (*Sturnus vulgaris*) and common kingfishers (*Alcedo atthis*) confirm part of this trend where larger clutches produce lighter hatchlings (Reid, Monaghan and Ruxton, 2000; Čech and Čech, 2017). What these studies do not confirm is that smaller clutches also produce lighter nestlings, as seen in our study. We could not find any studies showing a quadratic relationship between clutch size and hatch weights. The main reason for larger clutches producing smaller hatchings is that larger clutches cannot be uniformly incubated which leads to sub-optimal conditions for embryonic development compared to smaller clutches (Reid, Monaghan and Ruxton, 2000). However, this does not explain why smaller clutches resulted in smaller hatchlings in Navarino. It could be that smaller clutches are produced by poorer quality or older adults who may be less efficient in incubation resulting in poorer embryonic development that produce nestlings of lower quality. There is evidence for this theory where higher quality (more experienced/older) adults produced larger hatchlings (Lepage, Gauthier and Desrochers, 1998) and very old adults produce poorer quality hatchlings (Beamonte-Barrientos et al., 2010). There is also clear evidence that lower quality or very old adults in captivity and in the wild produce smaller clutch sizes and often lower quality hatchlings (Slagsvold and Lifjeld, 1990; Descamps et al., 2011). In Navarino, clutches of 1-3 eggs are uncommon (with clutches of 1-3 eggs making up only 14.8% of all clutch sizes). Thus, most clutches consisted of 4 eggs or more. This may mean that adults who produce small clutches may have pre-conditioned circumstances for doing so. For example, smaller clutches are often produced later in the season which is known to produce smaller nestlings (Crick, Gibbons and Magrath, 1993; Decker, Conway and Fontaine, 2012). Or adults may be older and senescence could be a reason for smaller clutches being produced that contain smaller nestlings. A study on blue footed boobies (*Sula nebouxii*) by Beamonte-Barrientos et al., (2010) found a quadratic effect of female age on egg production with old and young females producing smaller clutches on average, which may help to explain our finding. Clutch/brood size had no effect on hatch weights, growth rates or pre-fledging weights in Pucón.

#### Temporal changes of hatching and fledging weight and growth rates

We found that hatch weights are in decline in Navarino and Pucón since 2018. In Navarino, hatch weights have decreased on average 4% each year and hatch weights in Pucón have decreased 6% each year. However, despite the decline in hatch weights, growth rates and pre-fledgling weights are unchanged. This shows some level of mitigation, where parents likely trade-off more parental provisioning to mitigate the consequences of changing climatic conditions. If climatic conditions worsen, increased parental provisioning is not infinite and at some point, increased parental provisioning will not be able to mitigate small hatching weights and smaller nestlings by increased provisioning. It is likely that nestlings who are smaller at hatching receive more parental provisioning, this is likely because growth rates are unchanged, but hatch weights are in decline. This is particularly important in Navarino where winters are cold with heavy snow, and nestlings need to be large when they fledge to survive their first winter (see Botero-Delgadillo et al 2017).

#### Predation impacts on nestling development

Nest predation is present in Pucón and is a potentially newly emerging threat in Navarino. In Pucón the colocolo opossum (*Dromiciops gliroides*), an endangered marsupial, commonly invades nest boxes, eats the eggs and/or nestlings and use nest boxes to hibernate (Merino et al., 2009). The predation levels were not recorded and were more commonly on eggs than nestlings (Pers. Obs.). In addition, the predation rates of opossums on Rayaditos generally are not known. In Navarino, we have discovered a new potential predator of nestlings, the Magellanic woodpecker (*Campephilus magellanicus*) who has recently evolved the behaviour of using it’s arrow like tongue to spear nestlings through the nest box cavity hole, this happened in three nest boxes in 2023, 2/17 of which were occupied by Rayaditos and the third was occupied by the southern house wren (*Troglodytes musculus)*. In the case of the Rayadito occupied boxes, two nestlings from a brood of four were taken and the other, one nestling from a brood of five was taken. In the case of the southern house wren, all seven nestlings were taken and the nest box nearly completely dismantled by woodpecker attack. This is a new finding, and we do not know the impact (if any) on Rayadito fitness/behaviour.

Predation poses many challenges and its effect on nestling development is mixed. The theory is that under predation as a selection pressure; nestlings will grow faster and fledge earlier to avoid being predated in the nest. Hence, they should grow faster to reach optimum pre-fledging weight. One study by Tuero et al., (2018) on scissor-tailed flycatchers (*Tyrannus forficatus*) found no effect of predation to nestlings on nestling growth rates. This paper does not list which predators were present. More specifically, this study looked at populations with and without predation and growth rates were similar in both populations. However, the populations not only differed with respect to predation but also were located at different latitudes with different climatic variables, like our study sites. They found that rainfall was the key driver of growth rates, where wetter conditions resulted in higher growth rates, as rainfall is thought to increase insect abundances, the flycatcher’s food source. Typically, we would expect that higher predation risk would cause nestlings to grow faster in Pucón, but unless we measure the nestling period in both locations, we cannot know either way.

#### Concluding statements and recommendations for future research

Climate had an observable impact on nestling development in Patagonia. We have established that there is a potential trade-off from adults who provision nestlings more through extra feeding when hatchlings are smaller (because pre-fledging weights are unchanged between years and unaffected by climate/biotics factors). However, resources are limited and adults cannot exponentially provide parental care, as the climate continues to change for the worse, this will increase climatic selection pressures and result in increased intraspecific competition for food and territory. As the climate is expected to change dramatically over the next few decades (Natalia et al., 2020; Chemke, Ming and Yuval, 2022) we can only assume that this will intensify these selection pressures and present new challenges for this species and others like it. The impact of climate should be closely monitored, especially in the habitats we have studied that are sensitive to climate change. Future research should focus on testing the effects of climate in the field on nestling development, although it is not yet clear how this could be done. What could be done however, is a meta-analysis of the varying climatic and biotic impacts on nestling development across different species and different habitats. Collating a wide range of taxa in this way will allow us to better understand the mechanisms by which climate change is affecting nestling development and how this may shape the future of nestling development in birds.

## Supporting information

Supplementary data and R scripts

