## Supplementary data and R scripts for "Adaptation to climate driven environments in a Patagonian suboscine passerine": Climwin_Growth_Rates.R

```
library(ggplot2)
library(lme4)
library(multcompView)
library(multcomp)
library(glmmTMB)
library(DHARMa)
library(climwin)


NVENV <- read.csv("ENV_DATA_NVRN_ALL.csv", sep=",")        # NAVARINO WEATHER
PCENV <- read.csv("ENV_DATA_PCN_ALL.csv", sep=",")         # PUCON WEATHER
ENV <- read.csv("ENV_DATA_ALL_1823.csv", sep=",")          # ALL WEATHER DATA

OGRNV <- read.csv("OVERALL_GROWTH_NVRN.csv", sep=",")       # ALL OVERALL GROWTH NVRN
OGRPC <- read.csv("OVERALL_GROWTH_PCN.csv", sep=",")       # ALL OVERALL GROWTH PCN
OGRALL <- read.csv("OVERALL_GROWTH_ALL.csv", sep=",")       # ALL OVERALL GROWTH


D1NVRN <- read.csv("D1NVRN.csv", sep=",")                  # HATCH WEIGHTS CLIMWIN NVRN
D1PCN <- read.csv("D1PCN.csv", sep=",")                    # HATCH WEIGHTS CLIMWIN PCN
D1ALL <- read.csv("D1ALL.csv", sep=",")                    # HATCH WEIGHTS ALL

NVENV$LOCATION <- as.factor(NVENV$LOCATION)
NVENV$YEAR <- as.factor(NVENV$YEAR)
NVENV$MONTH <- as.factor(NVENV$MONTH)
NVENV$NEWDATE <- as.Date(NVENV$DATE,
                         format = "%d/%m/%Y")
NVENV$MONTHYEAR <- interaction(NVENV$MONTH, NVENV$YEAR)

PCENV$LOCATION <- as.factor(PCENV$LOCATION)
PCENV$YEAR <- as.factor(PCENV$YEAR)
PCENV$MONTH <- as.factor(PCENV$MONTH)
PCENV$NEWDATE <- as.Date(PCENV$DATE,
                         format = "%d/%m/%Y")
PCENV$MONTHYEAR <- interaction(PCENV$MONTH, PCENV$YEAR)

ENV$LOCATION <- as.factor(ENV$LOCATION)
ENV$YEAR <- as.factor(ENV$YEAR)
ENV$MONTH <- as.factor(ENV$MONTH)
ENV$NEWDATE <- as.Date(ENV$DATE,
                         format = "%d/%m/%Y")
ENV$MONTHYEAR <- interaction(ENV$MONTH, ENV$YEAR)


OGRNV$LOCATION <- as.factor(OGRNV$LOCATION)
OGRNV$YEAR <- as.factor(OGRNV$YEAR)
OGRNV$NEST_BOX <- as.factor(OGRNV$NEST_BOX)
OGRNV$BROOD <- as.factor(OGRNV$BROOD)
OGRNV$AGE <- as.factor(OGRNV$AGE)
OGRNV$MONTH <- as.factor(OGRNV$MONTH)
OGRNV$GROUP <- as.factor(OGRNV$GROUP)
OGRNV$NEWDATE <- as.Date(OGRNV$DATE,
                         format = "%d/%m/%Y")


OGRPC$LOCATION <- as.factor(OGRPC$LOCATION)
OGRPC$YEAR <- as.factor(OGRPC$YEAR)
OGRPC$NEST_BOX <- as.factor(OGRPC$NEST_BOX)
OGRPC$BROOD <- as.numeric(OGRPC$BROOD)
OGRPC$AGE <- as.factor(OGRPC$AGE)
OGRPC$MONTH <- as.factor(OGRPC$MONTH)
OGRPC$GROUP <- as.factor(OGRPC$GROUP)
OGRPC$NEWDATE <- as.Date(OGRPC$DATE,
                         format = "%d/%m/%Y")

OGRALL$LOCATION <- as.factor(OGRALL$LOCATION)
OGRALL$YEAR <- as.factor(OGRALL$YEAR)
OGRALL$NEST_BOX <- as.factor(OGRALL$NEST_BOX)
OGRALL$NEWDATE <- as.Date(OGRALL$DATE,
                         format = "%d/%m/%Y")
OGRALL$GROUP <- interaction(OGRALL$LOCATION, OGRALL$YEAR)


D1NVRN$LOCATION <- as.factor(D1NVRN$LOCATION)
D1NVRN$YEAR <- as.factor(D1NVRN$YEAR)
D1NVRN$NEST_BOX <- as.factor(D1NVRN$NEST_BOX)
D1NVRN$BROOD <- as.factor(D1NVRN$BROOD)
D1NVRN$AGE <- as.factor(D1NVRN$AGE)
D1NVRN$MONTH <- as.factor(D1NVRN$MONTH)
D1NVRN$NEWDATE <- as.Date(D1NVRN$DATE,
                          format = "%d/%m/%Y")
D1NVRN$GROUP <- interaction(D1NVRN$LOCATION, D1NVRN$YEAR)


D1PCN$LOCATION <- as.factor(D1PCN$LOCATION)
D1PCN$YEAR <- as.factor(D1PCN$YEAR)
D1PCN$NEST_BOX <- as.factor(D1PCN$NEST_BOX)
D1PCN$BROOD <- as.factor(D1PCN$BROOD)
D1PCN$AGE <- as.factor(D1PCN$AGE)
D1PCN$MONTH <- as.factor(D1PCN$MONTH)
D1PCN$NEWDATE <- as.Date(D1PCN$DATE,
                         format = "%d/%m/%Y")
D1PCN$GROUP <- interaction(D1PCN$LOCATION, D1PCN$YEAR)

D1ALL$LOCATION <- as.factor(D1ALL$LOCATION)
D1ALL$YEAR <- as.factor(D1ALL$YEAR)
D1ALL$NEST_BOX <- as.factor(D1ALL$NEST_BOX)
D1ALL$NEWDATE <- as.Date(D1ALL$DATE,
                          format = "%d/%m/%Y")
D1ALL$GROUP <- interaction(D1ALL$LOCATION, D1ALL$YEAR)
str(D1ALL)

########
#              NAVARINO HATCH WEIGHTS CLIMWIN
########


D1NVAVTEMP <- slidingwin(xvar = list(Temp = NVENV$Av_temp),
                         exclude = NA,
                         cdate = NVENV$NEWDATE,
                         bdate = D1NVRN$NEWDATE,
                         baseline = lmer(log10(BODY_MASS) ~ YEAR + (1|NEST_BOX), control=lmerControl(calc.derivs=F), data=D1NVRN),
                         cinterval = "day",
                         range = c(25, 0),
                         type = "relative",
                         stat = c("mean"),
                         func = c("lin", "quad"))

D1NVAVTEMP$combos
summary(D1NVAVTEMP[[1]]$BestModel)


lm1=lm(yvar~factor(YEAR)+climate, data=D1NVAVTEMP[[2]]$BestModelData)
lmer1=lmer(yvar~factor(YEAR)+climate+(1|NEST_BOX), data=D1NVAVTEMP[[2]]$BestModelData)
anova(lmer1,lm1)

BEST=D1NVAVTEMP[[1]]$BestModelData

summary(D1NVAVTEMP[[1]]$BestModel)

testDispersion(lmer1)
simulationOutput <- simulateResiduals(fittedModel =lmer1,re.form = ~0, plot = F)
simulate(lmer1,1000)
plot(simulationOutput)
testDispersion(simulationOutput)
res_sim <- DHARMa::simulateResiduals(D1NVAVTEMP[[1]]$BestModel, re.form = NULL)
plot(res_sim)

plotwin(dataset=D1NVAVTEMP[[1]]$Dataset)

BEST <- D1NVAVTEMP[[1]]$BestModelData

plot(D1NVAVTEMP[[1]]$BestModelData)

D1NVRN1 <- D1NVAVTEMP[[1]]$Dataset
summary(D1NVRN1)

RandGrow=randwin(repeats = 1000,  
                 window = "sliding", 
                 xvar=list(Temp = NVENV$Av_temp), 
                 cdate=NVENV$NEWDATE,
                 bdate = D1NVRN$NEWDATE,
                 baseline= lmer(log10(BODY_MASS) ~ YEAR + (1|NEST_BOX),control=lmerControl(calc.derivs=F), data=D1NVRN), 
                 cinterval = "day",
                 range=c(25,25),
                 type = "relative",
                 stat= c("mean"), 
                 func= c("lin"))

plotall(dataset = D1NVAVTEMP[[1]]$Dataset, datasetrand = RandGrow[[1]],
        cw1 = 0.95, cw2 = 0.5, cw3 = 0.25,
        title = NULL, arrow = FALSE)


############
#####                MIN TEMP AND NVRN HATCH WEIGHTS
############


D1NVMINTEMP <- slidingwin(xvar = list(Temp = NVENV$Min_Temp),
                          exclude = NA,
                          cdate = NVENV$NEWDATE,
                          bdate = D1NVRN$NEWDATE,
                          baseline = lmer(log10(BODY_MASS) ~ YEAR + (1|NEST_BOX), control=lmerControl(calc.derivs=F), data=D1NVRN), 
                          cinterval = "day",
                          range = c(25, 0),
                          type = "relative",
                          stat = c("mean"),
                          func = c("lin","quad"))

D1NVMINTEMP$combos
summary(D1NVMINTEMP[[1]]$BestModel)

lm1=lm(yvar~factor(YEAR)+climate, data=D1NVMINTEMP[[1]]$BestModelData)
lmer1=lmer(yvar~factor(YEAR)+climate+(1|NEST_BOX), data=D1NVMINTEMP[[1]]$BestModelData)
anova(lmer1,lm1)

simulationOutput <- simulateResiduals(fittedModel = D1NVMINTEMP[[1]]$BestModel, plot = F)
plot(simulationOutput)
testDispersion(simulationOutput)

res_sim <- DHARMa::simulateResiduals(D1NVMINTEMP[[1]]$BestModel, re.form = NULL)
plot(res_sim)

plotwin(dataset=D1NVMINTEMP[[1]]$Dataset)

BEST1 <- D1NVMINTEMP[[1]]$BestModelData

plot(D1NVMINTEMP[[1]]$BestModelData)

lm1=lm(yvar~YEAR+climate+I(climate^2), data=BEST1)
tuk=glht(lm1, linfct=mcp(YEAR="Tukey"))
plot(tuk)

ggplot(BEST1, aes(climate, yvar, colour=YEAR))+
  geom_point()+
  geom_line(aes(y=predict(lm1)), size=2)

RandGrow1=randwin(repeats = 1000, 
                  window = "sliding", 
                  xvar=list(Temp = NVENV$Min_Temp), 
                  cdate=NVENV$NEWDATE,
                  bdate = D1NVRN$NEWDATE,
                  cohort = D1NVRN$YEAR,
                  baseline= lmer(log10(BODY_MASS) ~ YEAR + (1|NEST_BOX),control=lmerControl(calc.derivs=F), data=D1NVRN), 
                  stat= "mean",
                  range=c(17,15), 
                  func="lin",
                  type ="relative", 
                  cinterval = "day")

plotall(dataset = D1NVMINTEMP[[1]]$Dataset, datasetrand = RandGrow1[[1]],
        cw1 = 0.95, cw2 = 0.5, cw3 = 0.25,
        title = NULL, arrow = FALSE)


############
#######             MAX TEMP AND NVRN HATCH WEIGHTS
############


D1NVMAXTEMP <- slidingwin(xvar = list(Temp = NVENV$Max_Temp),
                          exclude = NA,
                          cdate = NVENV$NEWDATE,
                          bdate = D1NVRN$NEWDATE,
                          baseline = lmer(log10(BODY_MASS) ~ YEAR + (1|NEST_BOX), control=lmerControl(calc.derivs=F), data=D1NVRN), 
                          cinterval = "day",
                          range = c(25, 0),
                          type = "relative", 
                          stat = c("mean"),
                          func = c("lin","quad"))

D1NVMAXTEMP$combos
summary(D1NVMAXTEMP[[1]]$BestModel)

lm1=lm(yvar~factor(YEAR)+climate, data=D1NVMAXTEMP[[1]]$BestModelData)
lmer1=lmer(yvar~factor(YEAR)+climate+(1|NEST_BOX), data=D1NVMAXTEMP[[1]]$BestModelData)
anova(lmer1,lm1)

simulationOutput <- simulateResiduals(fittedModel = D1NVMAXTEMP[[1]]$BestModel, plot = F)
plot(simulationOutput)
testDispersion(simulationOutput)

res_sim <- DHARMa::simulateResiduals(D1NVMAXTEMP[[1]]$BestModel, re.form = NULL)
plot(res_sim)

plotwin(dataset=D1NVMAXTEMP[[1]]$Dataset)

BEST2 <- D1NVMAXTEMP[[1]]$BestModelData

plot(D1NVMAXTEMP[[1]]$BestModelData)

D1NVRN1 <- D1NVMAXTEMP[[1]]$Dataset
summary(D1NVRN1)

RandGrow2=randwin(repeats = 1000, 
                  window = "sliding", 
                  xvar=list(Temp = NVENV$Max_Temp), 
                  cdate=NVENV$NEWDATE,
                  bdate = D1NVRN$NEWDATE,
                  cohort = D1NVRN$YEAR,
                  baseline= lmer(log10(BODY_MASS) ~ YEAR + (1|NEST_BOX), control=lmerControl(calc.derivs=F), data=D1NVRN), 
                  stat= "mean", 
                  range=c(25,25), 
                  func="lin",
                  type ="relative", 
                  cinterval = "day")

plotall(dataset = D1NVMAXTEMP[[1]]$Dataset, datasetrand = RandGrow2[[1]],
        cw1 = 0.95, cw2 = 0.5, cw3 = 0.25,
        title = NULL, arrow = FALSE)


###############
#########               RAIN AND NVRN HATCH WEIGHTS
###############


D1NVRAIN <- slidingwin(xvar = list(Rain = NVENV$Acc_mill),
                       exclude = NA,
                       cdate = NVENV$NEWDATE,
                       bdate = D1NVRN$NEWDATE,
                       baseline = lmer(log10(BODY_MASS) ~ YEAR + (1|NEST_BOX), control=lmerControl(calc.derivs=F), data=D1NVRN), 
                       cinterval = "day",
                       range = c(25, 0),
                       type = "relative", 
                       stat = c("mean"),
                       func = c("lin","quad"))

D1NVRAIN$combos
summary(D1NVRAIN[[2]]$BestModel)

lm1=lm(yvar~factor(YEAR)+climate+I(climate^2), data=D1NVRAIN[[2]]$BestModelData)
lmer1=lmer(yvar~factor(YEAR)+climate+(1|NEST_BOX), data=D1NVRAIN[[2]]$BestModelData)
anova(lmer1,lm1)

simulationOutput <- simulateResiduals(fittedModel = D1NVRAIN[[2]]$BestModel, plot = F)
plot(simulationOutput)
testDispersion(simulationOutput)

res_sim <- DHARMa::simulateResiduals(D1NVRAIN[[2]]$BestModel, re.form = NULL)
plot(res_sim)

plotwin(dataset=D1NVRAIN[[2]]$Dataset)

BEST3 <- D1NVRAIN[[2]]$BestModelData

plot(D1NVRAIN[[2]]$BestModelData)

D1NVRN1 <- D1NVRAIN[[2]]$Dataset
summary(D1NVRN1)

RandGrow=randwin(repeats = 1000, 
                 window = "sliding", 
                 xvar=list(Rain = NVENV$Acc_mill), 
                 cdate=NVENV$NEWDATE,
                 bdate = D1NVRN$NEWDATE,
                 cohort = D1NVRN$YEAR,
                 baseline= lmer(log10(BODY_MASS) ~ YEAR + (1|NEST_BOX), control=lmerControl(calc.derivs=F), data=D1NVRN), 
                 stat= "mean",
                 range=c(23,7), 
                 func="quad",
                 type ="relative", 
                 cinterval = "day")

plotall(dataset = D1NVRAIN[[2]]$Dataset, datasetrand = RandGrow[[1]],
        cw1 = 0.95, cw2 = 0.5, cw3 = 0.25,
        title = NULL, arrow = FALSE)

D1NVRN$RAIN_LIN=D1NVRAIN[[2]]$BestModelData$climate
D1NVRN$RAIN_QUAD=D1NVRAIN[[2]]$BestModelData$climate^2

############
######               WIND AND NVRN HATCH WEIGHTS
############

D1NVWIND <- slidingwin(xvar = list(Wind = NVENV$Av_ws_12),
                       exclude = NA,
                       cdate = NVENV$NEWDATE,
                       bdate = D1NVRN$NEWDATE,
                       baseline = lmer(log10(BODY_MASS) ~ YEAR + (1|NEST_BOX), control=lmerControl(calc.derivs=F), data=D1NVRN), 
                       cinterval = "day",
                       range = c(25, 0),
                       type = "relative", 
                       stat = c("mean"),
                       func = c("lin","quad"))

D1NVWIND$combos
summary(D1NVWIND[[1]]$BestModel)

lm1=lm(yvar~factor(YEAR)+climate, data=D1NVWIND[[1]]$BestModelData)
lmer1=lmer(yvar~factor(YEAR)+climate+(1|NEST_BOX), data=D1NVWIND[[1]]$BestModelData)
anova(lmer1,lm1)

simulationOutput <- simulateResiduals(fittedModel = D1NVWIND[[1]]$BestModel, plot = F)
plot(simulationOutput)
testDispersion(simulationOutput)

res_sim <- DHARMa::simulateResiduals(D1NVWIND[[1]]$BestModel, re.form = NULL)
plot(res_sim)

plotwin(dataset=D1NVWIND[[1]]$Dataset)

BEST4 <- D1NVWIND[[1]]$BestModelData

plot(D1NVWIND[[1]]$BestModelData)

D1NVRN1 <- D1NVWIND[[1]]$Dataset
summary(D1NVRN1)

RandGrow=randwin(repeats = 1000, 
                 window = "sliding", 
                 xvar=list(Wind = NVENV$Av_ws_12), 
                 cdate=NVENV$NEWDATE,
                 bdate = D1NVRN$NEWDATE,
                 cohort = D1NVRN$YEAR,
                 baseline= lmer(log10(BODY_MASS) ~ YEAR + (1|NEST_BOX), control=lmerControl(calc.derivs=F), data=D1NVRN), 
                 stat= "mean",
                 range=c(11,11), 
                 func="lin",
                 type ="relative", 
                 cinterval = "day")

plotall(dataset = D1NVWIND[[1]]$Dataset, datasetrand = RandGrow[[1]],
        cw1 = 0.95, cw2 = 0.5, cw3 = 0.25,
        title = NULL, arrow = FALSE)


#######
###         PCN HATCH WEIGHTS AND AVERAGE TEMPERATURE
#######

D1PCNAVTEMP <- slidingwin(xvar = list(Temp = PCENV$Av_temp),
                          exclude = NA,
                          cdate = PCENV$NEWDATE,
                          bdate = D1PCN$NEWDATE,
                          baseline = lmer(log10(BODY_MASS) ~ YEAR + (1|NEST_BOX), control=lmerControl(calc.derivs=F), data=D1PCN), 
                          cinterval = "day",
                          range = c(25, 0),
                          type = "relative", 
                          stat = c("mean"),
                          func = c("lin","quad"))

D1PCNAVTEMP$combos
summary(D1PCNAVTEMP[[1]]$BestModel)

lm1=lm(yvar~factor(YEAR)+climate, data=D1PCNAVTEMP[[1]]$BestModelData)
lmer1=lmer(yvar~factor(YEAR)+climate+(1|NEST_BOX), data=D1PCNAVTEMP[[1]]$BestModelData)
anova(lmer1,lm1)

simulationOutput <- simulateResiduals(fittedModel = D1PCNAVTEMP[[1]]$BestModel, plot = T)
plot(simulationOutput)
testDispersion(simulationOutput)

res_sim <- DHARMa::simulateResiduals(D1PCNAVTEMP[[1]]$BestModel, re.form = NULL)
plot(res_sim)

plotwin(dataset=D1PCNAVTEMP[[1]]$Dataset)

BEST5 <- D1PCNAVTEMP[[1]]$BestModelData

plot(D1PCNAVTEMP[[1]]$BestModelData)

D1PCN1 <- D1PCNAVTEMP[[1]]$Dataset
summary(D1PCN1)

RandGrow3=randwin(repeats = 1000, 
                  window = "sliding", 
                  xvar=list(Temp = PCENV$Av_temp), 
                  cdate=PCENV$NEWDATE,
                  bdate = D1PCN$NEWDATE,
                  cohort = D1PCN$YEAR,
                  baseline = lmer(log10(BODY_MASS) ~ YEAR + (1|NEST_BOX), control=lmerControl(calc.derivs=F), data=D1PCN), 
                  stat= "mean", 
                  range=c(20,19), 
                  func="lin",
                  type ="relative", 
                  cinterval = "day")

plotall(dataset = D1PCNAVTEMP[[1]]$Dataset, datasetrand = RandGrow3[[1]],
        cw1 = 0.95, cw2 = 0.5, cw3 = 0.25,
        title = NULL, arrow = FALSE)


#########
####           MIN TEMP HATCH WEIGHTS PCN
#########

D1PCNMINTEMP <- slidingwin(xvar = list(Temp = PCENV$Min_Temp),
                           exclude = NA,
                           cdate = PCENV$NEWDATE,
                           bdate = D1PCN$NEWDATE,
                           baseline = lmer(log10(BODY_MASS) ~ YEAR + (1|NEST_BOX), control=lmerControl(calc.derivs=F), data=D1PCN), 
                           cinterval = "day",
                           range = c(25, 0),
                           type = "relative", 
                           stat = c("mean"),
                           func = c("lin","quad"))

D1PCNMINTEMP$combos
summary(D1PCNMINTEMP[[1]]$BestModel)

lm1=lm(yvar~factor(YEAR)+climate, data=D1PCNMINTEMP[[1]]$BestModelData)
lmer1=lmer(yvar~factor(YEAR)+climate+(1|NEST_BOX), data=D1PCNMINTEMP[[1]]$BestModelData)
anova(lmer1,lm1)

simulationOutput <- simulateResiduals(fittedModel = D1PCNMINTEMP[[1]]$BestModel, plot = T)
plot(simulationOutput)
testDispersion(simulationOutput)

res_sim <- DHARMa::simulateResiduals(D1PCNMINTEMP[[1]]$BestModel, re.form = NULL)
plot(res_sim)

plotwin(dataset=D1PCNMINTEMP[[1]]$Dataset)

BEST6 <- D1PCNMINTEMP[[1]]$BestModelData

plot(D1PCNMINTEMP[[1]]$BestModelData)

D1PCN1 <- D1PCNMINTEMP[[1]]$Dataset
summary(D1PCN1)

RandGrow=randwin(repeats = 1000, 
                 window = "sliding", 
                 xvar=list(Temp = PCENV$Min_Temp), 
                 cdate=PCENV$NEWDATE,
                 bdate = D1PCN$NEWDATE,
                 cohort = D1PCN$YEAR,
                 baseline = lmer(log10(BODY_MASS) ~ YEAR + (1|NEST_BOX), control=lmerControl(calc.derivs=F), data=D1PCN), 
                 stat= "mean", 
                 range=c(19,18), 
                 func="lin",
                 type ="relative", 
                 cinterval = "day")

plotall(dataset = D1PCNMINTEMP[[1]]$Dataset, datasetrand = RandGrow[[1]],
        cw1 = 0.95, cw2 = 0.5, cw3 = 0.25,
        title = NULL, arrow = FALSE)

D1PCN$MIN_TEMP_LIN=D1PCNMINTEMP[[1]]$BestModelData$climate

###########
#####          MAX TEMP HATCH WEIGHTS PCN
###########


D1PCMAXTEMP <- slidingwin(xvar = list(Temp = PCENV$Max_Temp),
                          exclude = NA,
                          cdate = PCENV$NEWDATE,
                          bdate = D1PCN$NEWDATE,
                          baseline = lmer(log10(BODY_MASS) ~ YEAR + (1|NEST_BOX), control=lmerControl(calc.derivs=F), data=D1PCN), 
                          cinterval = "day",
                          range = c(25, 0),
                          type = "relative", 
                          stat = c("mean"),
                          func = c("lin","quad"))

D1PCMAXTEMP$combos
summary(D1PCMAXTEMP[[1]]$BestModel)

lm1=lm(yvar~factor(YEAR)+climate, data=D1PCMAXTEMP[[1]]$BestModelData)
lmer1=lmer(yvar~factor(YEAR)+climate+(1|NEST_BOX), data=D1PCMAXTEMP[[1]]$BestModelData)
anova(lmer1,lm1)


simulationOutput <- simulateResiduals(fittedModel = D1PCMAXTEMP[[1]]$BestModel, plot = T)
plot(simulationOutput)
testDispersion(simulationOutput)

res_sim <- DHARMa::simulateResiduals(D1PCMAXTEMP[[1]]$BestModel, re.form = NULL)
plot(res_sim)

plotwin(dataset=D1PCMAXTEMP[[1]]$Dataset)

BEST6 <- D1PCMAXTEMP[[1]]$BestModelData

plot(D1PCMAXTEMP[[1]]$BestModelData)

D1PCN1 <- D1PCMAXTEMP[[1]]$Dataset
summary(D1PCN1)

RandGrow=randwin(repeats = 1000, 
                 window = "sliding", 
                 xvar=list(Temp = PCENV$Max_Temp), 
                 cdate=PCENV$NEWDATE,
                 bdate = D1PCN$NEWDATE,
                 cohort = D1PCN$YEAR,
                 baseline = lmer(log10(BODY_MASS) ~ YEAR + (1|NEST_BOX), control=lmerControl(calc.derivs=F), data=D1PCN), 
                 stat= "mean", 
                 range=c(14,13), 
                 func="lin",
                 type ="relative", 
                 cinterval = "day")

plotall(dataset = D1PCMAXTEMP[[1]]$Dataset, datasetrand = RandGrow[[1]],
        cw1 = 0.95, cw2 = 0.5, cw3 = 0.25,
        title = NULL, arrow = FALSE)


############
####              RAIN HATCH WEIGHTS PCN
############


D1PCRAIN <- slidingwin(xvar = list(Rain = PCENV$Acc_mill),
                       exclude = NA,
                       cdate = PCENV$NEWDATE,
                       bdate = D1PCN$NEWDATE,
                       baseline = lmer(log10(BODY_MASS) ~ YEAR + (1|NEST_BOX), control=lmerControl(calc.derivs=F), data=D1PCN), 
                       cinterval = "day",
                       range = c(25, 0),
                       type = "relative", 
                       stat = c("mean"),
                       func = c("lin","quad"))

D1PCRAIN$combos
summary(D1PCRAIN[[1]]$BestModel)

lm1=lm(yvar~factor(YEAR)+climate, data=D1PCRAIN[[1]]$BestModelData)
lmer1=lmer(yvar~factor(YEAR)+climate+(1|NEST_BOX), data=D1PCRAIN[[1]]$BestModelData)
anova(lmer1,lm1)

simulationOutput <- simulateResiduals(fittedModel = D1PCRAIN[[1]]$BestModel, plot = T)
testDispersion(simulationOutput)

res_sim <- DHARMa::simulateResiduals(D1PCRAIN[[1]]$BestModel, re.form = NULL)
plot(res_sim)

plotwin(dataset=D1PCRAIN[[1]]$Dataset)

BEST7 <- D1PCRAIN[[1]]$BestModelData

plot(D1PCRAIN[[1]]$BestModelData)

D1PCN1 <- D1PCRAIN[[1]]$Dataset
summary(D1PCN1)

RandGrow=randwin(repeats = 1000, 
                 window = "sliding", 
                 xvar=list(Rain = PCENV$Acc_mill), 
                 cdate=PCENV$NEWDATE,
                 bdate = D1PCN$NEWDATE,
                 cohort = D1PCN$YEAR,
                 baseline = lmer(log10(BODY_MASS) ~ YEAR + (1|NEST_BOX), control=lmerControl(calc.derivs=F), data=D1PCN), 
                 stat= "mean", 
                 range=c(15,9), 
                 func="lin",
                 type ="relative", 
                 cinterval = "day")

plotall(dataset = D1PCRAIN[[1]]$Dataset, datasetrand = RandGrow[[1]],
        cw1 = 0.95, cw2 = 0.5, cw3 = 0.25,
        title = NULL, arrow = FALSE)

D1PCN$RAIN_LIN=D1PCRAIN[[1]]$BestModelData$climate

#############
######               WIND HATCH WEIGHTS PCN
############# 


D1PCWIND <- slidingwin(xvar = list(Wind = PCENV$Av_ws_12),
                       exclude = NA,
                       cdate = PCENV$NEWDATE,
                       bdate = D1PCN$NEWDATE,
                       baseline = lmer(log10(BODY_MASS) ~ YEAR + (1|NEST_BOX), control=lmerControl(calc.derivs=F), data=D1PCN), 
                       cinterval = "day",
                       range = c(25, 0),
                       type = "relative", 
                       stat = c("mean"),
                       func = c("lin","quad"))

D1PCWIND$combos
summary(D1PCWIND[[2]]$BestModel)

lm1=lm(yvar~factor(YEAR)+climate^2, data=D1PCRAIN[[2]]$BestModelData)
lmer1=lmer(yvar~factor(YEAR)+climate^2+(1|NEST_BOX), data=D1PCRAIN[[2]]$BestModelData)
anova(lmer1,lm1)

simulationOutput <- simulateResiduals(fittedModel = D1PCWIND[[2]]$BestModel, plot = T)
plot(simulationOutput) 
testDispersion(simulationOutput)

res_sim <- DHARMa::simulateResiduals(D1PCWIND[[2]]$BestModel, re.form = NULL)
plot(res_sim)


plotwin(dataset=D1PCWIND[[2]]$Dataset)

BEST8 <- D1PCWIND[[2]]$BestModelData

plot(D1PCWIND[[2]]$BestModelData)

D1PCN1 <- D1PCWIND[[2]]$Dataset
summary(D1PCN1)

RandGrow=randwin(repeats = 1000, 
                 window = "sliding", 
                 xvar=list(Wind = PCENV$Av_ws_12), 
                 cdate=PCENV$NEWDATE,
                 bdate = D1PCN$NEWDATE,
                 cohort = D1PCN$YEAR,
                 baseline = lmer(log10(BODY_MASS) ~ YEAR + (1|NEST_BOX), control=lmerControl(calc.derivs=F), data=D1PCN), 
                 stat= "mean", 
                 range=c(11,10), 
                 func="quad",
                 type ="relative", 
                 cinterval = "day")

plotall(dataset = D1PCWIND[[2]]$Dataset, datasetrand = RandGrow[[1]],
        cw1 = 0.95, cw2 = 0.5, cw3 = 0.25,
        title = NULL, arrow = FALSE)

D1PCN$WIND_LIN=D1PCWIND[[2]]$BestModelData$climate
D1PCN$WIND_QUAD=D1PCWIND[[2]]$BestModelData$climate^2

#######
######                 NAVARINO PRE-FLEDGING WEIGHTS
######


BODYMASSWIN <- slidingwin(xvar = list(Temp = NVENV$Av_temp),
                          exclude = NA,
                          cdate = NVENV$NEWDATE,
                          bdate = OGRNV$NEWDATE,
                          baseline = lm(log10(BODY_MASS) ~ YEAR, control=lmerControl(calc.derivs=F), data=OGRNV), 
                          cinterval = "day",
                          range = c(30,0),
                          type = "relative",
                          stat = c("mean"),
                          func = c("lin","quad"))

BODYMASSWIN$combos
summary(BODYMASSWIN[[1]]$BestModel)

lm1=lm(yvar~factor(YEAR)+climate, data=BODYMASSWIN[[1]]$BestModelData)
lmer1=lmer(yvar~factor(YEAR)+climate+(1|NEST_BOX), data=BODYMASSWIN[[1]]$BestModelData)
anova(lmer1,lm1)


simulationOutput <- simulateResiduals(fittedModel = BODYMASSWIN[[1]]$BestModel, plot = T)
plot(simulationOutput) 
testDispersion(simulationOutput)

res_sim <- DHARMa::simulateResiduals(BODYMASSWIN[[1]]$BestModel, re.form = NULL)
plot(res_sim)

plotwin(dataset=BODYMASSWIN[[1]]$Dataset)

BESTBM <- BODYMASSWIN[[1]]$BestModelData
plot(BESTBM$yvar, BESTBM$climate)


plot(BODYMASSWIN[[1]]$BestModelData)

PEAKAVTEMP <- BODYMASSWIN[[1]]$Dataset
summary(PEAKAVTEMP)

RandGrow=randwin(repeats = 1000, 
                 window = "sliding", 
                 xvar=list(Temp = NVENV$Av_temp), 
                 cdate=NVENV$NEWDATE,
                 bdate = OGRNV$NEWDATE,
                 cohort = OGRNV$YEAR,
                 baseline = lm(log10(BODY_MASS) ~ YEAR, control=lmerControl(calc.derivs=F), data=OGRNV), 
                 stat= "mean", 
                 range=c(27,27), 
                 func="lin",
                 type ="relative", 
                 cinterval = "day")

plotall(dataset = BODYMASSWIN[[1]]$Dataset, datasetrand = RandGrow[[1]],
        cw1 = 0.95, cw2 = 0.5, cw3 = 0.25,
        title = NULL, arrow = FALSE)


#########
#####               NVRN MAX TEMP AND D16 WEIGHT
##########


BODYMASSWINMAX <- slidingwin(xvar = list(Temp = NVENV$Max_Temp),
                             exclude = NA,
                             cdate = NVENV$NEWDATE,
                             bdate = OGRNV$NEWDATE,
                             baseline = lm(log10(BODY_MASS) ~ YEAR, control=lmerControl(calc.derivs=F), data = OGRNV), 
                             cinterval = "day",
                             range = c(30,0),
                             type = "relative",
                             stat = c("mean"),
                             func = c("lin","quad"))

BODYMASSWINMAX$combos
summary(BODYMASSWINMAX[[1]]$BestModel)

lm1=lm(yvar~factor(YEAR)+climate, data=BODYMASSWINMAX[[1]]$BestModelData)
lmer1=lmer(yvar~factor(YEAR)+climate+(1|NEST_BOX), data=BODYMASSWINMAX[[1]]$BestModelData)
anova(lmer1,lm1)


simulationOutput <- simulateResiduals(fittedModel = BODYMASSWINMAX[[1]]$BestModel, plot = T)
plot(simulationOutput) 
testDispersion(simulationOutput)

res_sim <- DHARMa::simulateResiduals(BODYMASSWINMAX[[1]]$BestModel, re.form = NULL)
plot(res_sim)

plotwin(dataset=BODYMASSWINMAX[[1]]$Dataset)

BESTBMMAX <- BODYMASSWINMAX[[1]]$BestModelData

plot(BODYMASSWINMAX[[1]]$BestModelData)

MAXTEMPD16W <- BODYMASSWINMAX[[1]]$Dataset
summary(MAXTEMPD16W)

RandGrow=randwin(repeats = 1000, 
                 window = "sliding", 
                 xvar=list(Temp = NVENV$Max_Temp), 
                 cdate=NVENV$NEWDATE,
                 bdate = OGRNV$NEWDATE,
                 cohort = OGRNV$YEAR,
                 baseline = lm(log10(BODY_MASS) ~ YEAR, control=lmerControl(calc.derivs=F), data = OGRNV), 
                 stat= "mean", 
                 range=c(20,18), 
                 func="lin",
                 type ="relative", 
                 cinterval = "day")

plotall(dataset = BODYMASSWINMAX[[1]]$Dataset, datasetrand = RandGrow[[1]],
        cw1 = 0.95, cw2 = 0.5, cw3 = 0.25,
        title = NULL, arrow = FALSE)


#########
#####            NVRN MIN TEMP
#########


BODYMASSWINMIN <- slidingwin(xvar = list(Temp = NVENV$Min_Temp),
                             exclude = NA,
                             cdate = NVENV$NEWDATE,
                             bdate = OGRNV$NEWDATE,
                             baseline = lm(log10(BODY_MASS) ~ YEAR, control=lmerControl(calc.derivs=F), data = OGRNV), 
                             cinterval = "day",
                             range = c(30,0),
                             type = "relative",
                             stat = c("mean"),
                             func = c("lin","quad"))

BODYMASSWINMIN$combos
summary(BODYMASSWINMIN[[1]]$BestModel)

lm1=lm(yvar~factor(YEAR)+climate, data=BODYMASSWINMIN[[1]]$BestModelData)
lmer1=lmer(yvar~factor(YEAR)+climate+(1|NEST_BOX), data=BODYMASSWINMIN[[1]]$BestModelData)
anova(lmer1,lm1)

simulationOutput <- simulateResiduals(fittedModel = BODYMASSWINMIN[[1]]$BestModel, plot = T)
plot(simulationOutput) 
testDispersion(simulationOutput)

res_sim <- DHARMa::simulateResiduals(BODYMASSWINMIN[[1]]$BestModel, re.form = NULL)
plot(res_sim)

plotwin(dataset=BODYMASSWINMIN[[1]]$Dataset)

BESTBMMIN <- BODYMASSWINMIN[[1]]$BestModelData
plot(BESTBMMIN$yvar, BESTBMMIN$climate)


plot(BODYMASSWINMIN[[1]]$BestModelData)

MINTEMPD16W <- BODYMASSWINMIN[[1]]$Dataset
summary(MINTEMPD16W)

RandGrow=randwin(repeats = 1000, 
                 window = "sliding", 
                 xvar=list(Temp = NVENV$Min_Temp), 
                 cdate=NVENV$NEWDATE,
                 bdate = OGRNV$NEWDATE,
                 cohort = OGRNV$YEAR,
                 baseline = lm(log10(BODY_MASS) ~ YEAR, control=lmerControl(calc.derivs=F), data = OGRNV), 
                 stat= "mean", 
                 range=c(30,30), 
                 func="lin",
                 type ="relative", 
                 cinterval = "day")

plotall(dataset = BODYMASSWINMIN[[1]]$Dataset, datasetrand = RandGrow[[1]],
        cw1 = 0.95, cw2 = 0.5, cw3 = 0.25,
        title = NULL, arrow = FALSE)


###########
######          NVRN RAINFALL AND D16 WEIGHT CLIMWIN
###########


BODYMASSRAIN <- slidingwin(xvar = list(Rain = NVENV$Acc_mill),
                           exclude = NA,
                           cdate = NVENV$NEWDATE,
                           bdate = OGRNV$NEWDATE,
                           baseline = lm(log10(BODY_MASS) ~ YEAR, control=lmerControl(calc.derivs=F), data = OGRNV), 
                           cinterval = "day",
                           range = c(30,0),
                           type = "relative",
                           stat = c("mean"),
                           func = c("lin","quad"))

BODYMASSRAIN$combos
summary(BODYMASSRAIN[[2]]$BestModel)

lm1=lm(yvar~factor(YEAR)+climate^2, data=BODYMASSRAIN[[2]]$BestModelData)
lmer1=lmer(yvar~factor(YEAR)+climate^2+(1|NEST_BOX), data=BODYMASSRAIN[[2]]$BestModelData)
anova(lmer1,lm1)


simulationOutput <- simulateResiduals(fittedModel = BODYMASSRAIN[[2]]$BestModel, plot = T)
plot(simulationOutput) 
testDispersion(simulationOutput)

res_sim <- DHARMa::simulateResiduals(BODYMASSRAIN[[2]]$BestModel, re.form = NULL)
plot(res_sim)

plotwin(dataset=BODYMASSRAIN[[2]]$Dataset)

BESTRAIN <- BODYMASSRAIN[[2]]$BestModelData
plot(BESTRAIN$yvar, BESTRAIN$climate)


plot(BODYMASSRAIN[[2]]$BestModelData)

RAIND16W <- BODYMASSRAIN[[2]]$Dataset
summary(RAIND16W)

RandGrow=randwin(repeats = 1000, 
                 window = "sliding", 
                 xvar=list(Rain = NVENV$Acc_mill), 
                 cdate=NVENV$NEWDATE,
                 bdate = OGRNV$NEWDATE,
                 cohort = OGRNV$YEAR,
                 baseline = lm(log10(BODY_MASS) ~ YEAR, control=lmerControl(calc.derivs=F), data = OGRNV), 
                 stat= "mean", 
                 range=c(28,26), 
                 func="quad",
                 type ="relative", 
                 cinterval = "day")

plotall(dataset = BODYMASSRAIN[[2]]$Dataset, datasetrand = RandGrow[[1]],
        cw1 = 0.95, cw2 = 0.5, cw3 = 0.25,
        title = NULL, arrow = FALSE)


############# 
#######                 NVRN D16 WEIGHT AND WIND SPEED
#############


BODYMASSWIND <- slidingwin(xvar = list(Wind = NVENV$Av_ws_12),
                           exclude = NA,
                           cdate = NVENV$NEWDATE,
                           bdate = OGRNV$NEWDATE,
                           baseline = lm(log10(BODY_MASS) ~ YEAR, control=lmerControl(calc.derivs=F), data = OGRNV), 
                           cinterval = "day",
                           range = c(30,0),
                           type = "relative",
                           stat = c("mean"),
                           func = c("lin","quad"))

BODYMASSWIND$combos
summary(BODYMASSWIND[[2]]$BestModel)

lm1=lm(yvar~factor(YEAR)+climate^2, data=BODYMASSWIND[[2]]$BestModelData)
lmer1=lmer(yvar~factor(YEAR)+climate^2+(1|NEST_BOX), data=BODYMASSWIND[[2]]$BestModelData)
anova(lmer1,lm1)

simulationOutput <- simulateResiduals(fittedModel = BODYMASSWIND[[2]]$BestModel, plot = T)
plot(simulationOutput) 
testDispersion(simulationOutput)

res_sim <- DHARMa::simulateResiduals(BODYMASSWIND[[2]]$BestModel, re.form = NULL)
plot(res_sim)

plotwin(dataset=BODYMASSWIND[[2]]$Dataset)

BESTWIND <- BODYMASSWIND[[2]]$BestModelData
plot(BESTWIND$yvar, BESTWIND$climate)

plot(BODYMASSWIND[[2]]$BestModelData)

WINDD16W <- BODYMASSWIND[[2]]$Dataset
summary(WINDD16W)

RandGrow=randwin(repeats = 1000, 
                 window = "sliding", 
                 xvar=list(Wind = NVENV$Av_ws_12), 
                 cdate=NVENV$NEWDATE,
                 bdate = OGRNV$NEWDATE,
                 cohort = OGRNV$YEAR,
                 baseline = lm(log10(BODY_MASS) ~ YEAR, control=lmerControl(calc.derivs=F), data = OGRNV), 
                 stat= "mean", 
                 range=c(19,19),
                 func="quad",
                 type ="relative", 
                 cinterval = "day")

plotall(dataset = BODYMASSWIND[[2]]$Dataset, datasetrand = RandGrow[[1]],
        cw1 = 0.95, cw2 = 0.5, cw3 = 0.25,
        title = NULL, arrow = FALSE)


######
##           PUCON PRE-FLEDGING WEIGHT AND AVERAGE TEMP
#######


BODYMASSWINPC <- slidingwin(xvar = list(Temp = PCENV$Av_temp),
                            exclude = NA,
                            cdate = PCENV$NEWDATE,
                            bdate = OGRPC$NEWDATE,
                            baseline = lm(log10(BODY_MASS) ~ YEAR, control=lmerControl(calc.derivs=F), data = OGRPC), 
                            cinterval = "day",
                            range = c(30,0),
                            type = "relative",
                            stat = c("mean"),
                            func = c("lin","quad"))

BODYMASSWINPC$combos
summary(BODYMASSWINPC[[2]]$BestModel)

lm1=lm(yvar~factor(YEAR)+climate, data=BODYMASSWINPC[[2]]$BestModelData)
lmer1=lmer(yvar~factor(YEAR)+climate+(1|NEST_BOX), data=BODYMASSWINPC[[2]]$BestModelData)
anova(lmer1,lm1)

simulationOutput <- simulateResiduals(fittedModel = BODYMASSWINPC[[2]]$BestModel, plot = T)
plot(simulationOutput) 
testDispersion(simulationOutput)

res_sim <- DHARMa::simulateResiduals(BODYMASSWINPC[[2]]$BestModel, re.form = NULL)
plot(res_sim)

plotwin(dataset=BODYMASSWINPC[[2]]$Dataset)

BESTBM <- BODYMASSWINPC[[2]]$BestModelData
plot(BESTBM$yvar, BESTBM$climate)


plot(BODYMASSWINPC[[2]]$BestModelData)

PEAKAVTEMP <- BODYMASSWINPC[[2]]$Dataset
summary(PEAKAVTEMP)

RandGrow=randwin(repeats = 1000, 
                 window = "sliding", 
                 xvar=list(Temp = PCENV$Av_temp), 
                 cdate=PCENV$NEWDATE,
                 bdate = OGRPC$NEWDATE,
                 cohort = OGRPC$YEAR,
                 baseline = lm(log10(BODY_MASS) ~ YEAR, control=lmerControl(calc.derivs=F), data = OGRPC), 
                 stat= "mean", 
                 range=c(26,25), 
                 func="quad",
                 type ="relative", 
                 cinterval = "day")

plotall(dataset = BODYMASSWINPC[[2]]$Dataset, datasetrand = RandGrow[[1]],
        cw1 = 0.95, cw2 = 0.5, cw3 = 0.25,
        title = NULL, arrow = FALSE)

######
#########            PCN MAX TEMP AND D16 WEIGHT
######


BODYMASSWINPCMAX <- slidingwin(xvar = list(Temp = PCENV$Max_Temp),
                               exclude = NA,
                               cdate = PCENV$NEWDATE,
                               bdate = OGRPC$NEWDATE,
                               baseline = lm(log10(BODY_MASS) ~ YEAR, control=lmerControl(calc.derivs=F), data = OGRPC), 
                               cinterval = "day",
                               range = c(30,0),
                               type = "relative",
                               stat = c("mean"),
                               func = c("lin","quad"))

BODYMASSWINPCMAX$combos
summary(BODYMASSWINPCMAX[[1]]$BestModel)

lm1=lm(yvar~factor(YEAR)+climate, data=BODYMASSWINPCMAX[[1]]$BestModelData)
lmer1=lmer(yvar~factor(YEAR)+climate+(1|NEST_BOX), data=BODYMASSWINPCMAX[[1]]$BestModelData)
anova(lmer1,lm1)

simulationOutput <- simulateResiduals(fittedModel = BODYMASSWINPCMAX[[1]]$BestModel, plot = T)
plot(simulationOutput) 
testDispersion(simulationOutput)

res_sim <- DHARMa::simulateResiduals(BODYMASSWINPCMAX[[1]]$BestModel, re.form = NULL)
plot(res_sim)

plotwin(dataset=BODYMASSWINPCMAX[[1]]$Dataset)

BESTBMMAX <- BODYMASSWINPCMAX[[1]]$BestModelData
plot(BESTBMMAX$yvar, BESTBMMAX$climate)


plot(BODYMASSWINPCMAX[[1]]$BestModelData)

MAXTEMPD16W <- BODYMASSWINPCMAX[[1]]$Dataset
summary(MAXTEMPD16W)

RandGrow=randwin(repeats = 1000, 
                 window = "sliding", 
                 xvar=list(Temp = PCENV$Max_Temp), 
                 cdate=PCENV$NEWDATE,
                 bdate = OGRPC$NEWDATE,
                 cohort = OGRPC$YEAR,
                 baseline = lm(log10(BODY_MASS) ~ YEAR, control=lmerControl(calc.derivs=F), data = OGRPC), 
                 stat= "mean", 
                 range=c(29,9), 
                 func="quad",
                 type ="relative", 
                 cinterval = "day")

plotall(dataset = BODYMASSWINPCMAX[[2]]$Dataset, datasetrand = RandGrow[[1]],
        cw1 = 0.95, cw2 = 0.5, cw3 = 0.25,
        title = NULL, arrow = FALSE)


########
###            PUCON MIN TEMP AND PRE-FLEDGING WEIGHT
##########


BODYMASSWINPCMIN <- slidingwin(xvar = list(Temp = PCENV$Min_Temp),
                               exclude = NA,
                               cdate = PCENV$NEWDATE,
                               bdate = OGRPC$NEWDATE,
                               baseline = lmer(log10(BODY_MASS) ~ YEAR + (1|NEST_BOX), control=lmerControl(calc.derivs=F), data = OGRPC), 
                               cinterval = "day",
                               range = c(30,0),
                               type = "relative",
                               stat = c("mean"),
                               func = c("lin","quad"))

BODYMASSWINPCMIN$combos
summary(BODYMASSWINPCMIN[[1]]$BestModel)

lm1=lm(yvar~factor(YEAR)+climate, data=BODYMASSWINPCMIN[[1]]$BestModelData)
lmer1=lmer(yvar~factor(YEAR)+climate+(1|NEST_BOX), data=BODYMASSWINPCMIN[[1]]$BestModelData)
anova(lmer1,lm1)

simulationOutput <- simulateResiduals(fittedModel = BODYMASSWINPCMIN[[1]]$BestModel, plot = T)
plot(simulationOutput) 
testDispersion(simulationOutput)

res_sim <- DHARMa::simulateResiduals(BODYMASSWINPCMIN[[1]]$BestModel, re.form = NULL)
plot(res_sim)

plotwin(dataset=BODYMASSWINPCMIN[[1]]$Dataset)

BESTBMMIN <- BODYMASSWINPCMIN[[1]]$BestModelData
plot(BESTBMMIN$yvar, BESTBMMIN$climate)


plot(BODYMASSWINPCMIN[[1]]$BestModelData)

MINTEMPD16W <- BODYMASSWINPCMIN[[1]]$Dataset
summary(MINTEMPD16W)

RandGrow=randwin(repeats = 1000, 
                 window = "sliding", 
                 xvar=list(Temp = PCENV$Min_Temp), 
                 cdate=PCENV$NEWDATE,
                 bdate = OGRPC$NEWDATE,
                 cohort = OGRPC$YEAR,
                 baseline = lmer(log10(BODY_MASS) ~ YEAR + (1|NEST_BOX), control=lmerControl(calc.derivs=F), data = OGRPC), 
                 stat= "mean", 
                 range=c(0,0), 
                 func="lin",
                 type ="relative", 
                 cinterval = "day")

plotall(dataset = BODYMASSWINPCMIN[[1]]$Dataset, datasetrand = RandGrow[[1]],
        cw1 = 0.95, cw2 = 0.5, cw3 = 0.25,
        title = NULL, arrow = FALSE)


#########
#####         PUCON RAIN AND PRE-FLEDGING WEIGHT
#########


BODYMASSRAIN <- slidingwin(xvar = list(Rain = PCENV$Acc_mill),
                           exclude = NA,
                           cdate = PCENV$NEWDATE,
                           bdate = OGRPC$NEWDATE,
                           baseline = lm(BODY_MASS ~ YEAR, control=lmerControl(calc.derivs=F), data = OGRPC), 
                           cinterval = "day",
                           range = c(30,0),
                           type = "relative",
                           stat = c("mean"),
                           func = c("lin","quad"))

BODYMASSRAIN$combos
summary(BODYMASSRAIN[[1]]$BestModel)

lm1=lm(yvar~factor(YEAR)+climate, data=BODYMASSRAIN[[1]]$BestModelData)
lmer1=lmer(yvar~factor(YEAR)+climate+(1|NEST_BOX), data=BODYMASSRAIN[[1]]$BestModelData)
anova(lmer1,lm1)

simulationOutput <- simulateResiduals(fittedModel = BODYMASSRAIN[[1]]$BestModel, plot = T)
plot(simulationOutput) 
testDispersion(simulationOutput)

res_sim <- DHARMa::simulateResiduals(BODYMASSRAIN[[1]]$BestModel, re.form = NULL)
plot(res_sim)

plotwin(dataset=BODYMASSRAIN[[1]]$Dataset)

BESTRAIN <- BODYMASSRAIN[[1]]$BestModelData
plot(BESTRAIN$yvar, BESTRAIN$climate)


plot(BODYMASSRAIN[[1]]$BestModelData)

RAIND16W <- BODYMASSRAIN[[1]]$Dataset
summary(RAIND16W)

RandGrow=randwin(repeats = 1000, 
                 window = "sliding", 
                 xvar=list(Rain = PCENV$Acc_mill), 
                 cdate=PCENV$NEWDATE,
                 bdate = OGRPC$NEWDATE,
                 cohort = OGRPC$YEAR,
                 baseline = lm(BODY_MASS ~ YEAR, control=lmerControl(calc.derivs=F), data = OGRPC), 
                 stat= "mean", 
                 range=c(18,17), 
                 func="lin",
                 type ="relative", 
                 cinterval = "day")

plotall(dataset = BODYMASSRAIN[[1]]$Dataset, datasetrand = RandGrow[[1]],
        cw1 = 0.95, cw2 = 0.5, cw3 = 0.25,
        title = NULL, arrow = FALSE)


#########
###            Wind speed and pre-fledging weight Pucon
#########


BODYMASSWINPCD <- slidingwin(xvar = list(Wind = PCENV$Av_ws_12),
                             exclude = NA,
                             cdate = PCENV$NEWDATE,
                             bdate = OGRPC$NEWDATE,
                             baseline = lm(log10(BODY_MASS) ~ YEAR, control=lmerControl(calc.derivs=F), data = OGRPC), 
                             cinterval = "day",
                             range = c(30,0),
                             type = "relative",
                             stat = c("mean"),
                             func = c("lin","quad"))

BODYMASSWINPCD$combos
summary(BODYMASSWINPCD[[1]]$BestModel)

lm1=lm(yvar~factor(YEAR)+climate, data=BODYMASSWINPCD[[1]]$BestModelData)
lmer1=lmer(yvar~factor(YEAR)+climate+(1|NEST_BOX), data=BODYMASSWINPCD[[1]]$BestModelData)
anova(lmer1,lm1)


simulationOutput <- simulateResiduals(fittedModel = BODYMASSWINPCD[[1]]$BestModel, plot = T)
plot(simulationOutput) 
testDispersion(simulationOutput)

res_sim <- DHARMa::simulateResiduals(BODYMASSWINPCD[[1]]$BestModel, re.form = NULL)
plot(res_sim)

plotwin(dataset=BODYMASSWINPCD[[1]]$Dataset)

BESTWIND <- BODYMASSWINPCD[[1]]$BestModelData
plot(BESTWIND$yvar, BESTWIND$climate)


plot(BODYMASSWINPCD[[1]]$BestModelData)

WINDD16W <- BODYMASSWINPCD[[1]]$Dataset
summary(WINDD16W)

RandGrow=randwin(repeats = 1000, 
                 window = "sliding", 
                 xvar=list(Wind = PCENV$Av_ws_12), 
                 cdate=PCENV$NEWDATE,
                 bdate = OGRPC$NEWDATE,
                 cohort = OGRPC$YEAR,
                 baseline = lm(log10(BODY_MASS) ~ YEAR, control=lmerControl(calc.derivs=F), data = OGRPC), 
                 stat= "mean", 
                 range=c(22,22), 
                 func="quad",
                 type ="relative", 
                 cinterval = "day")

plotall(dataset = BODYMASSWINPCD[[2]]$Dataset, datasetrand = RandGrow[[1]],
        cw1 = 0.95, cw2 = 0.5, cw3 = 0.25,
        title = NULL, arrow = FALSE)


############
#####                Navarino Growth Rates and av_temp
###########


GrowthWin <- slidingwin(xvar = list(Temp = NVENV$Av_temp),
                        exclude = NA,
                        cdate = NVENV$NEWDATE,
                        bdate = OGRNV$NEWDATE,
                        baseline = lm(log10(OVERALL_GROWTH_PD) ~ YEAR, control=lmerControl(calc.derivs=F), data = OGRNV), 
                        cinterval = "day",
                        range = c(30, 0),
                        type = "relative", 
                        stat = c("mean"),
                        func = c("lin","quad"))

GrowthWin$combos
summary(GrowthWin[[2]]$BestModel)

lm1=lm(yvar~factor(YEAR)+climate^2, data=GrowthWin[[2]]$BestModelData)
lmer1=lmer(yvar~factor(YEAR)+climate^2+(1|NEST_BOX), data=GrowthWin[[2]]$BestModelData)
anova(lmer1,lm1)

simulationOutput <- simulateResiduals(fittedModel = GrowthWin[[2]]$BestModel, plot = T)
testDispersion(simulationOutput)

res_sim <- DHARMa::simulateResiduals(GrowthWin[[2]]$BestModel, re.form = NULL)
plot(res_sim)

plotwin(dataset=GrowthWin[[2]]$Dataset)

BESTGROWTH <- GrowthWin[[2]]$BestModelData

plot(GrowthWin[[2]]$BestModelData)

OGRNV1 <- GrowthWin[[2]]$Dataset
summary(OGRNV1)

RandGrow=randwin(repeats = 1000, 
                 window = "sliding", 
                 xvar=list(Temp = NVENV$Av_temp), 
                 cdate=NVENV$NEWDATE,
                 bdate = OGRNV$NEWDATE,
                 cohort = OGRNV$YEAR,
                 baseline = lm(log10(OVERALL_GROWTH_PD) ~ YEAR, control=lmerControl(calc.derivs=F), data = OGRNV), 
                 stat= "mean", 
                 range=c(26,25), 
                 func="quad",
                 type ="relative", 
                 cinterval = "day")

plotall(dataset = GrowthWin[[2]]$Dataset, datasetrand = RandGrow[[1]],
        cw1 = 0.95, cw2 = 0.5, cw3 = 0.25,
        title = NULL, arrow = FALSE)

OGRNV$AV_TEMP_LIN=GrowthWin[[2]]$BestModelData$climate
OGRNV$AV_TEMP_QUAD=GrowthWin[[2]]$BestModelData$climate^2


####################
#                       OVERALL NESTLING GROWTH AND MIN TEMPERATURE NVRN
####################


MinTempWin <- slidingwin(xvar = list(Temp = NVENV$Min_Temp),
                         exclude = NA,
                         cdate = NVENV$NEWDATE,
                         bdate = OGRNV$NEWDATE,
                         baseline = lm(log10(OVERALL_GROWTH_PD) ~ YEAR, control=lmerControl(calc.derivs=F), data = OGRNV), 
                         cinterval = "day",
                         range = c(30, 0),
                         type = "relative", 
                         stat = c("mean"),
                         func = c("lin","quad"))

MinTempWin$combos
summary(MinTempWin[[1]]$BestModel)

lm1=lm(yvar~factor(YEAR)+climate^2, data=MinTempWin[[2]]$BestModelData)
lmer1=lmer(yvar~factor(YEAR)+climate^2+(1|NEST_BOX), data=MinTempWin[[2]]$BestModelData)
anova(lmer1,lm1)

simulationOutput <- simulateResiduals(fittedModel = MinTempWin[[2]]$BestModel, plot = T)
plot(simulationOutput) 
testDispersion(simulationOutput)

res_sim <- DHARMa::simulateResiduals(MinTempWin[[2]]$BestModel, re.form = NULL)
plot(res_sim)

plotwin(dataset=MinTempWin[[2]]$Dataset)

BESTGROWTH1 <- MinTempWin[[2]]$BestModelData

plot(MinTempWin[[2]]$BestModelData)

OGRNV1 <- MinTempWin[[2]]$Dataset
summary(OGRNV1)

RandGrow=randwin(repeats = 1000, 
                 window = "sliding", 
                 xvar=list(Temp = NVENV$Min_Temp), 
                 cdate=NVENV$NEWDATE,
                 bdate = OGRNV$NEWDATE,
                 cohort = OGRNV$YEAR,
                 baseline = lm(log10(OVERALL_GROWTH_PD) ~ YEAR, control=lmerControl(calc.derivs=F), data = OGRNV), 
                 stat= "mean",
                 range=c(27,22), 
                 func="quad",
                 type ="relative",
                 cinterval = "day")

plotall(dataset = MinTempWin[[2]]$Dataset, datasetrand = RandGrow[[1]],
        cw1 = 0.95, cw2 = 0.5, cw3 = 0.25,
        title = NULL, arrow = FALSE)


##################
#                      OVERALL NESTLING GROWTH AND MAX TEMPERATURE NVRN
###################


MaxTempWin <- slidingwin(xvar = list(Temp = NVENV$Max_Temp),
                         exclude = NA,
                         cdate = NVENV$NEWDATE,
                         bdate = OGRNV$NEWDATE,
                         baseline = lm(log10(OVERALL_GROWTH_PD) ~ YEAR, control=lmerControl(calc.derivs=F), data = OGRNV), 
                         cinterval = "day",
                         range = c(30, 0),
                         type = "relative", 
                         stat = c("mean"),
                         func = c("lin","quad"))

MaxTempWin$combos
summary(MaxTempWin[[2]]$BestModel)

lm1=lm(yvar~factor(YEAR)+climate^2, data=MaxTempWin[[2]]$BestModelData)
lmer1=lmer(yvar~factor(YEAR)+climate^2+(1|NEST_BOX), data=MaxTempWin[[2]]$BestModelData)
anova(lmer1,lm1)

simulationOutput <- simulateResiduals(fittedModel = MaxTempWin[[2]]$BestModel, plot = T)
plot(simulationOutput) 
testDispersion(simulationOutput)

res_sim <- DHARMa::simulateResiduals(MaxTempWin[[2]]$BestModel, re.form = NULL)
plot(res_sim)

plotwin(dataset=MaxTempWin[[2]]$Dataset)

BESTGROWTH2 <- MaxTempWin[[2]]$BestModelData

plot(MaxTempWin[[2]]$BestModelData)

OGRNV10 <- MaxTempWin[[2]]$Dataset
summary(OGRNV10)

RandGrow=randwin(repeats = 1000, 
                 window = "sliding", 
                 xvar=list(Temp = NVENV$Max_Temp), 
                 cdate=NVENV$NEWDATE,
                 bdate = OGRNV$NEWDATE,
                 cohort = OGRNV$YEAR,
                 baseline = lm(log10(OVERALL_GROWTH_PD) ~ YEAR, control=lmerControl(calc.derivs=F), data = OGRNV), 
                 stat= "mean", 
                 range=c(27,26), 
                 func="quad",
                 type ="relative", 
                 cinterval = "day")

plotall(dataset = MaxTempWin[[2]]$Dataset, datasetrand = RandGrow[[1]],
        cw1 = 0.95, cw2 = 0.5, cw3 = 0.25,
        title = NULL, arrow = FALSE)


####################
#                     OVERALL NESTLING GROWTH AND RAINFALL NVRN
####################


GROWTHWINRAIN <- slidingwin(xvar = list(Rain = NVENV$Acc_mill),
                            exclude = NA,
                            cdate = NVENV$NEWDATE,
                            bdate = OGRNV$NEWDATE,
                            baseline = lm(log10(OVERALL_GROWTH_PD) ~ YEAR, control=lmerControl(calc.derivs=F), data = OGRNV), 
                            cinterval = "day",
                            range = c(30, 0),
                            type = "relative", 
                            stat = c("mean"),
                            func = c("lin","quad"))

GROWTHWINRAIN$combos
summary(GROWTHWINRAIN[[2]]$BestModel)

lm1=lm(yvar~factor(YEAR)+climate^2, data=GROWTHWINRAIN[[2]]$BestModelData)
lmer1=lmer(yvar~factor(YEAR)+climate^2+(1|NEST_BOX), data=GROWTHWINRAIN[[2]]$BestModelData)
anova(lmer1,lm1)

simulationOutput <- simulateResiduals(fittedModel = GROWTHWINRAIN[[2]]$BestModel, plot = T)
plot(simulationOutput) 
testDispersion(simulationOutput)

res_sim <- DHARMa::simulateResiduals(GROWTHWINRAIN[[2]]$BestModel, re.form = NULL)
plot(res_sim)

plotwin(dataset=GROWTHWINRAIN[[2]]$Dataset)

BEST1 <- GROWTHWINRAIN[[2]]$BestModelData

plot(GROWTHWINRAIN[[2]]$BestModelData)

OGRNV2 <- GROWTHWINRAIN[[2]]$Dataset
summary(OGRNV2)

RandGrow1=randwin(repeats = 1000, 
                  window = "sliding", 
                  xvar=list(Rain = NVENV$Acc_mill), 
                  cdate=NVENV$NEWDATE,
                  bdate = OGRNV$NEWDATE,
                  cohort = OGRNV$YEAR,
                  baseline = lm(log10(OVERALL_GROWTH_PD) ~ YEAR, control=lmerControl(calc.derivs=F), data = OGRNV), 
                  stat= "mean", 
                  range=c(28,26), 
                  func="quad",
                  type ="relative", 
                  cinterval = "day")

plotall(dataset = GROWTHWINRAIN[[2]]$Dataset, datasetrand = RandGrow1[[1]],
        cw1 = 0.95, cw2 = 0.5, cw3 = 0.25,
        title = NULL, arrow = FALSE)


###################
#                      OVERALL NESTLING GROWTH AND WIND SPEED NVRN
###################


GROWTHWINWIND <- slidingwin(xvar = list(Wind = NVENV$Av_ws_12),
                            exclude = NA,
                            cdate = NVENV$NEWDATE,
                            bdate = OGRNV$NEWDATE,
                            baseline = lm(log10(OVERALL_GROWTH_PD) ~ YEAR, control=lmerControl(calc.derivs=F), data = OGRNV), 
                            cinterval = "day",
                            range = c(30, 0),
                            type = "relative", 
                            stat = c("mean"),
                            func = c("lin","quad"))

GROWTHWINWIND$combos
summary(GROWTHWINWIND[[2]]$BestModel)

lm1=lm(yvar~factor(YEAR)+climate, data=GROWTHWINWIND[[1]]$BestModelData)
lmer1=lmer(yvar~factor(YEAR)+climate+(1|NEST_BOX), data=GROWTHWINWIND[[1]]$BestModelData)
anova(lmer1,lm1)

simulationOutput <- simulateResiduals(fittedModel = GROWTHWINWIND[[2]]$BestModel, plot = T)
plot(simulationOutput) 
testDispersion(simulationOutput)

res_sim <- DHARMa::simulateResiduals(GROWTHWINWIND[[2]]$BestModel, re.form = NULL)
plot(res_sim)


plotwin(dataset=GROWTHWINWIND[[2]]$Dataset)

BESTGROWTH3 <- GROWTHWINWIND[[2]]$BestModelData

plot(GROWTHWINWIND[[2]]$BestModelData)

OGRNV3 <- GROWTHWINWIND[[2]]$Dataset
summary(OGRNV3)

RandGrow2=randwin(repeats = 1000, 
                  window = "sliding", 
                  xvar=list(Wind = NVENV$Av_ws_12), 
                  cdate=NVENV$NEWDATE,
                  bdate = OGRNV$NEWDATE,
                  cohort = OGRNV$YEAR,
                  baseline = lm(log10(OVERALL_GROWTH_PD) ~ YEAR, control=lmerControl(calc.derivs=F), data = OGRNV), 
                  stat= "mean", 
                  range=c(19,19), 
                  func="quad",
                  type ="relative", 
                  cinterval = "day")

plotall(dataset = GROWTHWINWIND[[2]]$Dataset, datasetrand = RandGrow2[[1]],
        cw1 = 0.95, cw2 = 0.5, cw3 = 0.25,
        title = NULL, arrow = FALSE)


########
#######             PUCON OVERALL GROWTH RATES AND AVERAGE TEMP CLIMWIN
#########


PCTEMPGPD <- slidingwin(xvar = list(Temp = PCENV$Av_temp),
                        exclude = NA,
                        cdate = PCENV$NEWDATE,
                        bdate = OGRPC$NEWDATE,
                        baseline = lm(log10(OVERALL_GROWTH_PD) ~ YEAR, control=lmerControl(calc.derivs=F), data = OGRPC), 
                        cinterval = "day",
                        range = c(30,0),
                        type = "relative",
                        stat = c("mean"),
                        func = c("lin","quad"))

PCTEMPGPD$combos
summary(PCTEMPGPD[[2]]$BestModel)

lm1=lm(yvar~factor(YEAR)+climate^2, data=PCTEMPGPD[[2]]$BestModelData)
lmer1=lmer(yvar~factor(YEAR)+climate^2+(1|NEST_BOX), data=PCTEMPGPD[[2]]$BestModelData)
anova(lmer1,lm1)

simulationOutput <- simulateResiduals(fittedModel = PCTEMPGPD[[2]]$BestModel, plot = T)
plot(simulationOutput) 
testDispersion(simulationOutput)

res_sim <- DHARMa::simulateResiduals(PCTEMPGPD[[2]]$BestModel, re.form = NULL)
plot(res_sim)

plotwin(dataset=PCTEMPGPD[[2]]$Dataset)

BEST <- PCTEMPGPD[[2]]$BestModelData

plot(PCTEMPGPD[[2]]$BestModelData)

OGRPC1 <- PCTEMPGPD[[2]]$Dataset
summary(OGRPC1)

RandGrow=randwin(repeats = 1000, 
                 window = "sliding", 
                 xvar=list(Temp = PCENV$Av_temp), 
                 cdate=PCENV$NEWDATE,
                 bdate = OGRPC$NEWDATE,
                 cohort = OGRPC$YEAR,
                 baseline = lm(log10(OVERALL_GROWTH_PD) ~ YEAR, control=lmerControl(calc.derivs=F), data = OGRPC), 
                 stat= "mean", 
                 range=c(11,11), 
                 func="quad",
                 type ="relative", 
                 cinterval = "day")

plotall(dataset = PCTEMPGPD[[2]]$Dataset, datasetrand = RandGrow[[1]],
        cw1 = 0.95, cw2 = 0.5, cw3 = 0.25,
        title = NULL, arrow = FALSE)


####
#           OVERALL GROWTH RATES AND MAX TEMPERATURE PCN
####


PCMAXTEMP <- slidingwin(xvar = list(Temp = PCENV$Max_Temp),
                        exclude = NA,
                        cdate = PCENV$NEWDATE,
                        bdate = OGRPC$NEWDATE,
                        baseline = lmer(log10(OVERALL_GROWTH_PD) ~ YEAR + (1|NEST_BOX), control=lmerControl(calc.derivs=F), data = OGRPC), 
                        cinterval = "day",
                        range = c(30,0),
                        type = "relative",
                        stat = c("mean"),
                        func = c("lin","quad"))

PCMAXTEMP$combos
summary(PCMAXTEMP[[1]]$BestModel)

lm1=lm(yvar~factor(YEAR)+climate, data=PCMAXTEMP[[1]]$BestModelData)
lmer1=lmer(yvar~factor(YEAR)+climate+(1|NEST_BOX), data=PCMAXTEMP[[1]]$BestModelData)
anova(lmer1,lm1)

simulationOutput <- simulateResiduals(fittedModel = PCMAXTEMP[[1]]$BestModel, plot = T)
plot(simulationOutput) 
testDispersion(simulationOutput)

res_sim <- DHARMa::simulateResiduals(PCMAXTEMP[[1]]$BestModel, re.form = NULL)
plot(res_sim)

plotwin(dataset=PCMAXTEMP[[1]]$Dataset)

BEST1 <- PCMAXTEMP[[1]]$BestModelData

plot(PCMAXTEMP[[1]]$BestModelData)

OGRPC2 <- PCMAXTEMP[[1]]$Dataset
summary(OGRPC2)

RandGrow=randwin(repeats = 1000, 
                 window = "sliding", 
                 xvar=list(Temp = PCENV$Max_Temp), 
                 cdate=PCENV$NEWDATE,
                 bdate = OGRPC$NEWDATE,
                 cohort = OGRPC$YEAR,
                 baseline = lmer(log10(OVERALL_GROWTH_PD) ~ YEAR + (1|NEST_BOX), control=lmerControl(calc.derivs=F), data = OGRPC), 
                 stat= "mean", 
                 range=c(22,2), 
                 func="lin",
                 type ="relative", 
                 cinterval = "day")

plotall(dataset = PCMAXTEMP[[1]]$Dataset, datasetrand = RandGrow[[1]],
        cw1 = 0.95, cw2 = 0.5, cw3 = 0.25,
        title = NULL, arrow = FALSE)


###########
####              Growth rates and minimum temp Pucon
###########


PCMINTEMP <- slidingwin(xvar = list(Temp = PCENV$Min_Temp),
                        exclude = NA,
                        cdate = PCENV$NEWDATE,
                        bdate = OGRPC$NEWDATE,
                        baseline = lm(log10(OVERALL_GROWTH_PD) ~ YEAR, control=lmerControl(calc.derivs=F), data = OGRPC), 
                        cinterval = "day",
                        range = c(30,0),
                        type = "relative",
                        stat = c("mean"),
                        func = c("lin","quad"))

PCMINTEMP$combos
summary(PCMINTEMP[[1]]$BestModel)

lm1=lm(yvar~factor(YEAR)+climate^2, data=PCMINTEMP[[2]]$BestModelData)
lmer1=lmer(yvar~factor(YEAR)+climate^2+(1|NEST_BOX), data=PCMINTEMP[[2]]$BestModelData)
anova(lmer1,lm1)

simulationOutput <- simulateResiduals(fittedModel = PCMINTEMP[[2]]$BestModel, plot = T)
plot(simulationOutput) 
testDispersion(simulationOutput)

res_sim <- DHARMa::simulateResiduals(PCMINTEMP[[2]]$BestModel, re.form = NULL)
plot(res_sim)


plotwin(dataset=PCMINTEMP[[2]]$Dataset)

BEST <- PCMINTEMP[[2]]$BestModelData

plot(PCMINTEMP[[2]$BestModelData)

OGRPC1 <- PCMINTEMP[[2]]$Dataset
summary(OGRPC1)

RandGrow=randwin(repeats = 1000, 
                 window = "sliding", 
                 xvar=list(Temp = PCENV$Min_Temp), 
                 cdate=PCENV$NEWDATE,
                 bdate = OGRPC$NEWDATE,
                 cohort = OGRPC$YEAR,
                 baseline = lm(log10(OVERALL_GROWTH_PD) ~ YEAR, control=lmerControl(calc.derivs=F), data = OGRPC), 
                 stat= "mean", 
                 range=c(3,3), 
                 func="quad",
                 type ="relative", 
                 cinterval = "day")

plotall(dataset = PCMINTEMP[[2]]$Dataset, datasetrand = RandGrow[[1]],
        cw1 = 0.95, cw2 = 0.5, cw3 = 0.25,
        title = NULL, arrow = FALSE)


###
##            OVERALL GROWTH RATES AND RAIN PCN
###


PCRAIN <- slidingwin(xvar = list(Rain = PCENV$Acc_mill),
                     exclude = NA,
                     cdate = PCENV$NEWDATE,
                     bdate = OGRPC$NEWDATE,
                     baseline = lm(OVERALL_GROWTH_PD ~ YEAR, control=lmerControl(calc.derivs=F), data = OGRPC), 
                     cinterval = "day",
                     range = c(30,0),
                     type = "relative",
                     stat = c("mean"),
                     func = c("lin","quad"))

PCRAIN$combos
summary(PCRAIN[[2]]$BestModel)

lm1=lm(yvar~factor(YEAR)+climate^2, data=PCRAIN[[2]]$BestModelData)
lmer1=lmer(yvar~factor(YEAR)+climate^2+(1|NEST_BOX), data=PCRAIN[[2]]$BestModelData)
anova(lmer1,lm1)

simulationOutput <- simulateResiduals(fittedModel = PCRAIN[[2]]$BestModel, plot = T)
plot(simulationOutput) 
testDispersion(simulationOutput)

res_sim <- DHARMa::simulateResiduals(PCRAIN[[2]]$BestModel, re.form = NULL)
plot(res_sim)

plotwin(dataset=PCRAIN[[2]]$Dataset)

BEST <- PCRAIN[[2]]$BestModelData

plot(PCRAIN[[2]]$BestModelData)

OGRPC2 <- PCRAIN[[2]]$Dataset
summary(OGRPC2)

RandGrow=randwin(repeats = 1000, 
                 window = "sliding", 
                 xvar=list(Rain = PCENV$Acc_mill), 
                 cdate=PCENV$NEWDATE,
                 bdate = OGRPC$NEWDATE,
                 cohort = OGRPC$YEAR,
                 baseline = lm(OVERALL_GROWTH_PD ~ YEAR, control=lmerControl(calc.derivs=F), data = OGRPC), 
                 stat= "mean", 
                 range=c(8,2), 
                 func="quad",
                 type ="relative", 
                 cinterval = "day")

plotall(dataset = PCRAIN[[2]]$Dataset, datasetrand = RandGrow[[1]],
        cw1 = 0.95, cw2 = 0.5, cw3 = 0.25,
        title = NULL, arrow = FALSE)


###
####            OVERALL GROWTH RATES AND WIND SPEEDS PCN
###


PCWIND <- slidingwin(xvar = list(Wind = PCENV$Av_ws_12),
                     exclude = NA,
                     cdate = PCENV$NEWDATE,
                     bdate = OGRPC$NEWDATE,
                     baseline = lmer(log10(OVERALL_GROWTH_PD) ~ YEAR + (1|NEST_BOX), control=lmerControl(calc.derivs=F), data = OGRPC), 
                     cinterval = "day",
                     range = c(30,0),
                     type = "relative",
                     stat = c("mean"),
                     func = c("lin","quad"))

PCWIND$combos
summary(PCWIND[[2]]$BestModel)

lm1=lm(yvar~factor(YEAR)+climate^2, data=PCWIND[[2]]$BestModelData)
lmer1=lmer(yvar~factor(YEAR)+climate^2+(1|NEST_BOX), data=PCWIND[[2]]$BestModelData)
anova(lmer1,lm1)

simulationOutput <- simulateResiduals(fittedModel = PCWIND[[2]]$BestModel, plot = T)
plot(simulationOutput) 
testDispersion(simulationOutput)

res_sim <- DHARMa::simulateResiduals(PCWIND[[2]]$BestModel, re.form = NULL)
plot(res_sim)

plotwin(dataset=PCWIND[[2]]$Dataset)

BEST <- PCWIND[[2]]$BestModelData

plot(PCWIND[[2]]$BestModelData)

OGRPC1 <- PCWIND[[2]]$Dataset
summary(OGRPC1)

RandGrow=randwin(repeats = 1000,
                 window = "sliding", 
                 xvar=list(Wind = PCENV$Av_ws_12), 
                 cdate=PCENV$NEWDATE,
                 bdate = OGRPC$NEWDATE,
                 cohort = OGRPC$YEAR,
                 baseline = lmer(log10(OVERALL_GROWTH_PD) ~ YEAR + (1|NEST_BOX), control=lmerControl(calc.derivs=F), data = OGRPC), 
                 stat= "mean", 
                 range=c(27,4), 
                 func="lin",
                 type ="relative", 
                 cinterval = "day")

plotall(dataset = PCWIND[[1]]$Dataset, datasetrand = RandGrow[[1]],
        cw1 = 0.95, cw2 = 0.5, cw3 = 0.25,
        title = NULL, arrow = FALSE)


########                                                          ######################
###                       ALL HATCH WEIGHTS CLIMWIN               ############
########                                                       #############################

ALLWIND <- slidingwin(xvar = list(Wind = ENV$Av_ws_12),
                     exclude = NA,
                     cdate = ENV$NEWDATE,
                     bdate = D1ALL$NEWDATE,
                     baseline = lmer(log10(BODY_MASS) ~ YEAR + (1|NEST_BOX), control=lmerControl(calc.derivs=F), data = D1ALL), 
                     cinterval = "day",
                     range = c(30,0),
                     type = "relative",
                     stat = c("mean"),
                     func = c("lin","quad"), spatial = list(D1ALL$LOCATION, ENV$LOCATION))

ALLWIND$combos
summary(ALLWIND[[2]]$BestModel)

res_sim <- DHARMa::simulateResiduals(ALLWIND[[2]]$BestModel, re.form = NULL)
plot(res_sim)

plotwin(dataset=ALLWIND[[2]]$Dataset)

BEST <- ALLWIND[[2]]$BestModelData

plot(ALLWIND[[2]]$BestModelData)

D1ALL1 <- ALLWIND[[2]]$Dataset
summary(D1ALL1)

RandGrow=randwin(repeats = 1000,
                 window = "sliding", 
                 xvar=list(Wind = ENV$Av_ws_12), 
                 cdate=ENV$NEWDATE,
                 bdate = D1ALL$NEWDATE,
                 cohort = D1ALL$YEAR,
                 baseline = lmer(log10(BODY_MASS) ~ YEAR + (1|NEST_BOX), control=lmerControl(calc.derivs=F), data = D1ALL), 
                 stat= "mean", 
                 range=c(30,0), 
                 func="lin",
                 type ="relative", 
                 cinterval = "day") spatial = list(D1ALL$LOCATION, ENV$LOCATION)

plotall(dataset = ALLWIND[[1]]$Dataset, datasetrand = RandGrow[[1]],
        cw1 = 0.95, cw2 = 0.5, cw3 = 0.25,
        title = NULL, arrow = FALSE)

plot(ALLWIND[[1]]$BestModelData$climate)
```
