## Supplementary data and R scripts for "Adaptation to climate driven environments in a Patagonian suboscine passerine": Nestling_Growth_Code.R

```
library(MuMIn)
library(PerformanceAnalytics)
library(Hmisc)
library(lme4)
library(car)
library(pcr)
library(caret)
library(pls)
library(mgcv)
library(FactoMineR)
library(dplyr)
library(factoextra)
library(missMDA)
library(mgcViz)
library(emmeans)
library(visreg)
library(glmmTMB)
library(mgcv)
library(DHARMa)
library(psych)
library(ggeffects)
library(viridis)

NVENV <- read.csv("ENV_DATA_NVRN_ALL.csv", sep=",")        # NAVARINO WEATHER
PCENV <- read.csv("ENV_DATA_PCN_ALL.csv", sep=",")         # PUCON WEATHER

OGRNV <- read.csv("OVERALL_GROWTH_NVRN.csv", sep=",")       # ALL OVERALL GROWTH NVRN
OGRPC <- read.csv("OVERALL_GROWTH_PCN.csv", sep=",")       # ALL OVERALL GROWTH PCN

D1NVRN <- read.csv("D1NVRN.csv", sep=",")                  # HATCH WEIGHTS NVRN
D1PCN <- read.csv("D1PCN.csv", sep=",")                    # HATCH WEIGHTS PCN

D1ALL <- read.csv("D1ALL.csv", sep=",")                    # HATCH WEIGHTS ALL
NESTALL <- read.csv("NESTLING_GROWTH_PER_DAY.csv", sep=",") # ALL NESTLINGS GROWTH AND D16 BODY MASS


NVENV$LOCATION <- as.factor(NVENV$LOCATION)
NVENV$YEAR <- as.factor(NVENV$YEAR)
NVENV$MONTH <- as.factor(NVENV$MONTH)
NVENV$NEWDATE <- as.Date(NVENV$DATE,
                         format = "%d/%m/%Y")
NVENV$MONTHYEAR <- interaction(NVENV$MONTH, NVENV$YEAR)

PCENV$LOCATION <- as.factor(PCENV$LOCATION)
PCENV$YEAR <- as.factor(PCENV$YEAR)
PCENV$MONTH <- as.factor(PCENV$MONTH)
PCENV$NEWDATE <- as.Date(PCENV$DATE,
                         format = "%d/%m/%Y")
PCENV$MONTHYEAR <- interaction(PCENV$MONTH, PCENV$YEAR)


OGRNV$LOCATION <- as.factor(OGRNV$LOCATION)
OGRNV$YEAR <- as.factor(OGRNV$YEAR)
OGRNV$NEST_BOX <- as.factor(OGRNV$NEST_BOX)
OGRNV$BROOD <- as.numeric(OGRNV$BROOD)
OGRNV$AGE <- as.factor(OGRNV$AGE)
OGRNV$MONTH <- as.factor(OGRNV$MONTH)
OGRNV$GROUP <- as.factor(OGRNV$GROUP)
OGRNV$NEWDATE <- as.Date(OGRNV$DATE,
                         format = "%d/%m/%Y")
OGRNV$NB_YEAR <- paste(OGRNV$NEST_BOX, OGRNV$YEAR, sep = "_")
OGRNV$NB_YEAR <- as.factor(OGRNV$NB_YEAR)
OGRNV$YEAR <- factor(OGRNV$YEAR,
                     levels = c(1819, 1920, 2023),
                     labels = c("2018", "2019", "2023"))

OGRPC$LOCATION <- as.factor(OGRPC$LOCATION)
OGRPC$YEAR <- as.factor(OGRPC$YEAR)
OGRPC$NEST_BOX <- as.factor(OGRPC$NEST_BOX)
OGRPC$BROOD <- as.numeric(OGRPC$BROOD)
OGRPC$AGE <- as.factor(OGRPC$AGE)
OGRPC$MONTH <- as.factor(OGRPC$MONTH)
OGRPC$GROUP <- as.factor(OGRPC$GROUP)
OGRPC$NEWDATE <- as.Date(OGRPC$DATE,
                         format = "%d/%m/%Y")
OGRPC$NB_YEAR <- paste(OGRPC$NEST_BOX, OGRPC$YEAR, sep = "_")
OGRPC$NB_YEAR <- as.factor(OGRPC$NB_YEAR)


D1NVRN$YEAR <- factor(D1NVRN$YEAR,
                      levels = c(1819, 1920, 2023),
                      labels = c("2018", "2019", "2023"))
D1NVRN$LOCATION <- as.factor(D1NVRN$LOCATION)
D1NVRN$YEAR <- as.factor(D1NVRN$YEAR)
D1NVRN$NEST_BOX <- as.factor(D1NVRN$NEST_BOX)
D1NVRN$BROOD <- as.numeric(D1NVRN$BROOD)
D1NVRN$AGE <- as.factor(D1NVRN$AGE)
D1NVRN$MONTH <- as.factor(D1NVRN$MONTH)
D1NVRN$DAYS_TO_HATCH <- as.numeric(D1NVRN$DAYS_TO_HATCH)
D1NVRN$NEWDATE <- as.Date(D1NVRN$DATE,
                          format = "%d/%m/%Y")
D1NVRN$GROUP <- interaction(D1NVRN$LOCATION, D1NVRN$YEAR)
D1NVRN$NB_YEAR <- interaction(D1NVRN$NEST_BOX, D1NVRN$YEAR)
D1NVRN$NB_YEAR <- as.factor(D1NVRN$NB_YEAR)

D1PCN$YEAR <- factor(D1PCN$YEAR,
                     levels = c(1819, 1920, 2023),
                     labels = c("2018", "2019", "2023"))
D1PCN$LOCATION <- as.factor(D1PCN$LOCATION)
D1PCN$YEAR <- as.factor(D1PCN$YEAR)
D1PCN$NEST_BOX <- as.factor(D1PCN$NEST_BOX)
D1PCN$BROOD <- as.numeric(D1PCN$BROOD)
D1PCN$AGE <- as.factor(D1PCN$AGE)
D1PCN$MONTH <- as.factor(D1PCN$MONTH)
D1PCN$NEWDATE <- as.Date(D1PCN$DATE,
                         format = "%d/%m/%Y")
D1PCN$GROUP <- interaction(D1PCN$LOCATION, D1PCN$YEAR)
D1PCN$NB_YEAR <- interaction(D1PCN$NEST_BOX, D1PCN$YEAR)
D1PCN$NB_YEAR <- as.factor(D1PCN$NB_YEAR)


################
##                     Analysis of all nestlings
#################

str(D1ALL)
D1ALL$GROUP <- interaction(D1ALL$LOCATION, D1ALL$YEAR)

D1ALL$LOCATION <- as.factor(D1ALL$LOCATION)
D1ALL$YEAR <- as.factor(D1ALL$YEAR)
D1ALL$GROUP <- as.factor(D1ALL$GROUP)

str(NESTALL)

NESTALL$GROUP <- interaction(NESTALL$LOCATION, NESTALL$YEAR)
NESTALL$NB_YEAR <- interaction(NESTALL$NEST_BOX, NESTALL$YEAR)
NESTALL$NB_YEAR <- as.factor(NESTALL$NB_YEAR)

length(unique(NESTALL$NB_YEAR))

AM1 <- lm(BODY_MASS ~ LOCATION, data=D1ALL)
summary(AM1)

mean1 <- mean(D1PCN$BODY_MASS, na.rm = TRUE)
mean2 <- mean(D1NVRN$BODY_MASS, na.rm = TRUE)
percent_change <- ((mean2 - mean1) / mean1) * 100
percent_change


AN1 <- lm(OVERALL_GROWTH_PD ~ LOCATION, data=NESTALL)
summary(AN1)
mean3 <- mean(OGRNV$OVERALL_GROWTH_PD, na.rm = TRUE)
mean4 <- mean(OGRPC$OVERALL_GROWTH_PD, na.rm = TRUE)
percent_change <- ((mean3 - mean4) / mean3) * 100
percent_change

mean5 <- mean(OGRNV$BODY_MASS, na.rm = TRUE)
mean6 <- mean(OGRPC$BODY_MASS, na.rm = TRUE)
percent_change <- ((mean5 - mean6) / mean5) * 100
percent_change

AN2 <- lm(BODY_MASS ~ LOCATION, data=NESTALL)
summary(AN2)

mean(D1NVRN$BROOD)

##########
######           Modelling hatch weights in Navarino
##########

str(D1NVRN)
qqPlot(D1NVRN$BODY_MASS)
shapiro.test(D1NVRN$BODY_MASS)
hist(D1NVRN$BODY_MASS)
table(D1NVRN$BROOD)
table(D1NVRN$YEAR)

mean(D1NVRN$BODY_MASS)
sd(D1NVRN$BODY_MASS)


sapply(D1NVRN, function(x) sum(is.na(x) | is.nan(x) | is.infinite(x)))
# Removing rows with missing data in relevant columns
D1NVRN <- D1NVRN[complete.cases(D1NVRN[, c("BODY_MASS", "RAIN_LIN", "RAIN_QUAD", "DAYS_TO_HATCH", "BROOD", "NEST_BOX", "YEAR")]), ]

### first make some plots to see what the data looks like
plot(BODY_MASS~as.numeric(DAYS_TO_HATCH), data=D1NVRN)
plot(BODY_MASS~RAIN_LIN, data=D1NVRN)
plot(BODY_MASS~RAIN_QUAD, data=D1NVRN)
plot(BODY_MASS~YEAR, data=D1NVRN) # Looks like they're getting smaller
plot(BODY_MASS~as.numeric(BROOD), data=D1NVRN)# there may be a quadtatic relationship between brood and body mass - add brood^2 into models
plot(BODY_MASS~as.numeric(NEST_BOX), data=D1NVRN)

pairs(D1NVRN[, c("BODY_MASS", "INSECT_T","RAIN_QUAD","RAIN_LIN", "DAYS_TO_HATCH", "BROOD", "NEST_BOX", "YEAR")])

mean_by_year <- aggregate(BODY_MASS ~ YEAR,
                          data = D1NVRN,
                          FUN = mean,
                          na.rm = TRUE)

mean_by_year

mean_by_year$percent_change <- c(
  NA,
  diff(mean_by_year$BODY_MASS) / head(mean_by_year$BODY_MASS, -1) * 100
)

mean_by_year # 4% decrease each year

# full model
str(D1NVRN)
D1NVRN$BROOD <- as.numeric(D1NVRN$BROOD)

#D1NVRN$RAIN_centered <- scale(D1NVRN$RAIN_LIN, scale = FALSE)
#D1NVRN$RAIN_LIN <- D1NVRN$RAIN_centered
#D1NVRN$RAIN_QUAD <- D1NVRN$RAIN_centered^2

lmd1nv <- lmer(log10(BODY_MASS) ~ DAYS_TO_HATCH + YEAR + INSECT_T + BROOD + RAIN_LIN + RAIN_QUAD +
                 (1|NB_YEAR), data=D1NVRN)

vif(lmd1nv)
summary(lmd1nv)
drop1(lmd1nv, test="Chisq");
Sim=simulateResiduals(lmd1nv, re.form = ~0, plot = T)
testDispersion(Sim)

plot(BODY_MASS~RAIN_QUAD, data=D1NVRN)
plot(BODY_MASS~RAIN_LIN, data=D1NVRN)
plot(BODY_MASS ~ BROOD, data=D1NVRN)
plot(BODY_MASS ~ DAYS_TO_HATCH, data=D1NVRN)

D1NVRN$BROOD <- factor(D1NVRN$BROOD)
bsnv <- lm(BODY_MASS ~ BROOD, data=D1NVRN)
summary(bsnv)
emm <- emmeans(bsnv, ~ BROOD)
emm

######   RAINFALL QUAD   ##########
scale_x_continuous(breaks = seq(1, 3, by = 0.5))

ggplot(D1NVRN, aes(x = RAIN_LIN, y = BODY_MASS, color = as.factor(YEAR))) +
  geom_point(alpha = 0.6, size = 3) +
  geom_smooth(aes(group = YEAR),
              method = "lm",
              formula = y ~ poly(x, 2),
              se = FALSE,
              color = "black",
              linewidth = 1.3) +
  geom_smooth(method = "lm",
              formula = y ~ poly(x, 2),
              se = TRUE,
              linewidth = 1.5,
              alpha = 0.25) +
  scale_x_continuous(breaks = seq(1, 3, by = 0.5)) +
  scale_color_brewer(palette = "Set2", name = "Year") +
  labs(x = "Daily Total Rainfall (ml)", y = "Hatch Weight (g)") +
  theme_classic(base_size = 30) +
  theme(
    axis.title = element_text(face = "bold"),
    panel.border = element_rect(color = "black", fill = NA)
  )

#########     brood size

D1NVRN$BROOD <- factor(D1NVRN$BROOD)

ggplot(D1NVRN, aes(x = BROOD, y = BODY_MASS)) +
  geom_boxplot(
    width = 0.6,
    outlier.shape = NA,
    fill = "#4C72B0",
    alpha = 0.9
  ) +
  geom_jitter(
    width = 0.15,
    size = 2,
    alpha = 0.6,
    color = "black"
  ) +
  labs(
    title = "",
    x = "Clutch Size",
    y = "Hatching Weight (g)"
  ) +
  theme_classic(base_size = 14) +
  theme(
    plot.title = element_text(
      hjust = 0.5,
      face = "bold",
      size = 16
    ),
    axis.title = element_text(size = 24, color = "black",face = "bold"),
    axis.text = element_text(size = 22, color = "black"),
    axis.line = element_line(linewidth = 0.8),
    axis.ticks = element_line(linewidth = 0.8)
  )

table(D1NVRN$BROOD)

D1NVRN$BROOD_num <- as.numeric(as.character(D1NVRN$BROOD))

percent_1_3 <- sum(D1NVRN$BROOD %in% 1:3, na.rm = TRUE) /
  sum(!is.na(D1NVRN$BROOD)) * 100

percent_1_3

##########                                             #################
######           Modelling hatch weights in Pucon      #############
##########                                             ##################

str(D1PCN)

mean(D1PCN$BODY_MASS)
sd(D1PCN$BODY_MASS)

mean(D1PCN$BROOD)
table(D1PCN$YEAR)

plot(BODY_MASS~YEAR, data=D1PCN) # Hatch weights look smaller in 2019 than 2018
plot(BODY_MASS~RAIN_LIN, data=D1PCN)
plot(BODY_MASS~MIN_TEMP_LIN, data=D1PCN)
plot(BODY_MASS~WIND_LIN, data=D1PCN)
plot(BODY_MASS~WIND_QUAD, data=D1PCN)
plot(BODY_MASS~as.numeric(BROOD), data=D1PCN)# A bit quadratic but mostly linear - body mass decreases as broods get bigger
plot(BODY_MASS~as.numeric(NEST_BOX), data=D1PCN)

# MIN TEMP IS THE BEST HERE (LOWEST AIC) - but I do want to consider all weather so PCA is the best approach
AICc(lmer(BODY_MASS~INSECT_T+(1|NEST_BOX), data=D1PCN))
AICc(lmer(BODY_MASS~MIN_TEMP_LIN+(1|NEST_BOX), data=D1PCN))
AICc(lmer(BODY_MASS~RAIN_LIN+(1|NEST_BOX), data=D1PCN))
AICc(lmer(BODY_MASS~WIND_LIN+WIND_QUAD+(1|NEST_BOX), data=D1PCN))

mean_by_year <- aggregate(BODY_MASS ~ YEAR,
                          data = D1PCN,
                          FUN = mean,
                          na.rm = TRUE)

mean_by_year

mean_by_year$percent_change <- c(
  NA,
  diff(mean_by_year$BODY_MASS) / head(mean_by_year$BODY_MASS, -1) * 100
)

mean_by_year # 6% decrease each year

# Final model

#D1PCN$WIND_centered <- scale(D1PCN$WIND_LIN, scale = FALSE)
#D1PCN$WIND_LIN <- D1PCN$WIND_centered
#D1PCN$WIND_QUAD <- D1PCN$WIND_centered^2

lmhpc1 <- lmer(log10(BODY_MASS) ~ MIN_TEMP_LIN + RAIN_LIN + WIND_LIN + WIND_QUAD + INSECT_T + YEAR + BROOD + (1|NB_YEAR), data=D1PCN)

vif(lmhpc1) # All good
summary(lmhpc1)
drop1(lmhpc1, test="Chisq")
Sim=simulateResiduals(lmhpc1, re.form = ~0, plot = T)
testDispersion(Sim)

############       RAINFALL      ##############

ggplot(D1PCN, aes(x = RAIN_LIN,
                  y = BODY_MASS,
                  colour = as.factor(YEAR))) +
  geom_point(alpha = 0.8, size = 3) +
  geom_smooth(aes(group = 1),
              method = "lm",
              formula = y ~ x,
              se = TRUE,
              color = "black",
              fill = "gray70") +
  scale_color_manual(
    values = c("#8B0000", "#FF8C00", "#00008B", "#008B8B", "#654321"),
    name = "Year"
  ) +
  labs(x = "Total Daily Rainfall (ml)",
       y = "Hatch Weight (g)") +
  theme_classic(base_size = 28) +
  theme(
    axis.title = element_text(face = "bold"),
    panel.border = element_rect(color = "black", fill = NA),
    legend.position = "right"
  )

ggplot(D1PCN, aes(x = RAIN_LIN, y = BODY_MASS, color = as.factor(YEAR))) +
  geom_point(alpha = 0.8, size = 3) +
  geom_smooth(data = subset(D1PCN, YEAR %in% c(2018, 2019)),
              aes(group = YEAR),
              method = "lm",
              formula = y ~ x,
              se = FALSE,
              color = "black",
              linewidth = 2.2) +
  geom_smooth(data = subset(D1PCN, YEAR %in% c(2018, 2019)),
              aes(group = YEAR),
              method = "lm",
              formula = y ~ x,
              se = TRUE,
              linewidth = 1.5) +
  scale_color_manual(values = c("2018" = "#8B0000",  # dark red
                                "2019" = "#FF8C00"), # dark orange
                     name = "Year") +
  labs(x = "Total Daily Rainfall (ml)",
       y = "Hatch Weight (g)") +
  theme_classic(base_size = 28) +
  theme(
    axis.title = element_text(face = "bold"),
    panel.border = element_rect(color = "black", fill = NA),
    legend.position = "right"
  )


table(D1PCN$YEAR)
######    WIND SPEED 2    ###########

ggplot(D1PCN, aes(x = WIND_LIN,
                  y = BODY_MASS,
                  colour = as.factor(YEAR))) +
  geom_point(alpha = 0.8, size = 3) +
  
  geom_smooth(data = subset(D1PCN, YEAR %in% c(2018, 2019)),
              aes(group = YEAR),
              method = "lm",
              formula = y ~ poly(x, 2),
              se = TRUE,
              colour = NA,
              fill = "grey70",
              alpha = 0.4) +
  geom_smooth(data = subset(D1PCN, YEAR %in% c(2018, 2019)),
              aes(colour = as.factor(YEAR)),
              method = "lm",
              formula = y ~ poly(x, 2),
              se = FALSE,
              linewidth = 2) +
  
  scale_color_manual(
    values = c("2018" = "#1B263B",
               "2019" = "#7D3C98"),
    name = "Year"
  ) +
  labs(x = "Daily Average Wind Speed (km/h)", 
       y = "Hatch Weight (g)") +
  theme_classic(base_size = 28) +
  theme(
    axis.title = element_text(face = "bold"),
    panel.border = element_rect(color = "black", fill = NA),
    legend.position = "right"
  )


####        YEAR     #######

ggplot(D1PCN, aes(x = as.factor(YEAR), y = BODY_MASS, fill = as.factor(YEAR))) +
  geom_boxplot(color = "black") +
  scale_fill_manual(
    values = c("1819" = "#87CEFA",  # light sky blue
               "1920" = "#FFA07A"), # light salmon
    name = "Year"
  ) +
  scale_x_discrete(labels = c("1819" = "2019", "1920" = "2020")) +
  labs(x = "Year", 
       y = "Body Mass (g)") +
  theme_classic(base_size = 28) +
  theme(
    axis.title = element_text(face = "bold"),
    panel.border = element_rect(color = "black", fill = NA),
    legend.position = "right"
  )

colSums(is.na(D1PCN[, c("BODY_MASS", "MIN_TEMP_LIN", "RAIN_LIN", "WIND_QUAD", "BROOD", "INSECT_T", "YEAR")]))
sapply(D1PCN, function(x) sum(is.na(x) | is.nan(x) | is.infinite(x)))

summary(D1PCN[, c("BODY_MASS", "MIN_TEMP_LIN", "RAIN_LIN", "WIND_QUAD", "BROOD", "INSECT_T", "YEAR")])


############
###########        Growth rates in Navarino
############

str(OGRNV)
hist(OGRNV$OVERALL_GROWTH_PD)
qqPlot(OGRNV$OVERALL_GROWTH_PD)

mean(OGRNV$OVERALL_GROWTH_PD)
sd(OGRNV$OVERALL_GROWTH_PD)

mean(OGRNV$BODY_MASS)
sd(OGRNV$BODY_MASS)

table(OGRNV$YEAR)

plot(OVERALL_GROWTH_PD~INSECT_T, data=OGRNV)
plot(OVERALL_GROWTH_PD~AV_TEMP_QUAD, data=OGRNV)
plot(OVERALL_GROWTH_PD~DAYS_TO_HATCH, data=OGRNV)
plot(OVERALL_GROWTH_PD~BROOD, data=OGRNV) # Looks quadratic

AICc(lmer(OVERALL_GROWTH_PD~INSECT_T+(1|NEST_BOX), data=OGRNV))
AICc(lmer(OVERALL_GROWTH_PD~AV_TEMP_QUAD+(1|NEST_BOX), data=OGRNV))
AICc(lmer(OVERALL_GROWTH_PD~DAYS_TO_HATCH + (1|NEST_BOX), data=OGRNV))


sapply(OGRNV, function(x) sum(is.na(x) | is.nan(x) | is.infinite(x)))

OGRNV <- OGRNV[complete.cases(OGRNV[, c("OVERALL_GROWTH_PD", "AV_TEMP_QUAD","AV_TEMP_LIN", "DAYS_TO_HATCH", "BROOD", "NEST_BOX", "YEAR", "INSECT_T")]), ]

gr1 <- lm(OVERALL_GROWTH_PD ~ YEAR, data=OGRNV) # no change in growth rate over years
summary(gr1)

############# Full model

OGRNV$AV_TEMP_centered <- scale(OGRNV$AV_TEMP_LIN, scale = FALSE)
OGRNV$AV_TEMP_LIN <- OGRNV$AV_TEMP_centered
OGRNV$AV_TEMP_QUAD <- OGRNV$AV_TEMP_centered^2


lmd1nvg <- lmer(log10(OVERALL_GROWTH_PD) ~ AV_TEMP_LIN + AV_TEMP_QUAD +
                  YEAR + BROOD + INSECT_T + DAYS_TO_HATCH + (1|NB_YEAR), data=OGRNV)

vif(lmd1nvg) # all good
summary(lmd1nvg);
drop1(lmd1nvg, test="Chisq");  # Only Av temp quad
Sim=simulateResiduals(lmd1nvg, re.form = ~0, plot = T) # All good
testDispersion(Sim)

plot(OVERALL_GROWTH_PD ~ AV_TEMP_LIN, data=OGRNV)
plot(OVERALL_GROWTH_PD ~ AV_TEMP_QUAD, data=OGRNV)

### AVERAGE TEMP QUADRATIC  ########


ggplot(OGRNV, aes(x = AV_TEMP_LIN, y = OVERALL_GROWTH_PD, color = as.factor(YEAR))) +
  geom_point(alpha = 0.8, size = 3) +
  geom_smooth(aes(group = YEAR), method = "lm", formula = y ~ poly(x, 2), se = FALSE, color = "black", linewidth = 1.3) +
  geom_smooth(method = "lm", formula = y ~ poly(x, 2), se = TRUE, linewidth = 1.0, alpha = 0.25) +
  scale_color_manual(values = c("#1B263B", "#D35400", "#196F3D", "#7F4E1E", "#00008B"), name = "Year") +
  labs(x = "Daily Average Ambient Temperature (°C)", y = "Overall Growth per day (g)") +
  theme_classic(base_size = 22) +
  theme(axis.title = element_text(face = "bold"), panel.border = element_rect(color = "black", fill = NA), legend.position = "right")


################
###########                 Growth rates in Pucon
################


# No CLIMWIN results so other predictors only

str(OGRPC)
hist(OGRPC$OVERALL_GROWTH_PD)
table(OGRPC$BROOD)
table(OGRPC$YEAR)

sapply(OGRPC, function(x) sum(is.na(x) | is.nan(x) | is.infinite(x)))
# Data is clean

mean(OGRPC$OVERALL_GROWTH_PD)
sd(OGRPC$OVERALL_GROWTH_PD)

mean(OGRPC$BODY_MASS)
sd(OGRPC$BODY_MASS)


GRPCLM <- lmer(OVERALL_GROWTH_PD ~ YEAR + INSECT_T + BROOD + (1|NB_YEAR), data=OGRPC)
vif(GRPCLM)
summary(GRPCLM)
drop1(GRPCLM, test="Chisq");
Sim=simulateResiduals(GRPCLM, re.form = ~0, plot = T)
testDispersion(Sim)

###################
##########                   Pre-fledgling weights Navarino
###################

#CIMWIN found no weather correlations

sapply(OGRNV, function(x) sum(is.na(x) | is.nan(x) | is.infinite(x)))
colSums(is.na(OGRNV[, c("BODY_MASS", "DAYS_TO_HATCH", "INSECT_T", "YEAR")]))
OGRNV <- OGRNV[!is.na(OGRNV$DAYS_TO_HATCH), ]
str(OGRNV)

PFWNV <- lmer(log10(BODY_MASS) ~ YEAR + INSECT_T + BROOD + DAYS_TO_HATCH + (1|NB_YEAR), data=OGRNV)
summary(PFWNV)
drop1(PFWNV, test="Chisq");
vif(PFWNV) # ALL BELOW 4
Sim=simulateResiduals(PFWNV, re.form = ~0, plot = T) # all good
testDispersion(Sim)
plot(INSECT_T~BODY_MASS, data=OGRNV)

plot(BODY_MASS~INSECT_T, data=OGRNV)
plot(BODY_MASS~BROOD, data=OGRNV)
plot(BODY_MASS~DAYS_TO_HATCH, data=OGRNV)

AICc(lmer(BODY_MASS~INSECT_T+(1|NEST_BOX), data=OGRNV))
AICc(lmer(BODY_MASS~BROOD+(1|NEST_BOX), data=OGRNV))
AICc(lmer(BODY_MASS~DAYS_TO_HATCH + (1|NEST_BOX), data=OGRNV))
AICc(lmer(BODY_MASS~YEAR+ (1|NEST_BOX), data=OGRNV))

# To include hatch weight and growth rates as predictors

sapply(D1NVRN, function(x) sum(is.na(x) | is.nan(x) | is.infinite(x))) # 34 nestlings died before ringing so have no ring number
D1NVRN<- D1NVRN[!is.na(D1NVRN$RING), ] # Removed the 34 dead nestlings


OGRNV <- OGRNV %>%
  left_join(
    D1NVRN %>% select(RING, BODY_MASS),
    by = "RING"
  ) %>%
  rename(HATCH_WEIGHT = BODY_MASS.y)

PFWNV1 <- lmer(log10(BODY_MASS.x) ~ YEAR + INSECT_T + BROOD + DAYS_TO_HATCH + OVERALL_GROWTH_PD + (1|NB_YEAR), data=OGRNV)
summary(PFWNV1)
drop1(PFWNV1, test="Chisq");
vif(PFWNV1) # ALL BELOW 4
Sim=simulateResiduals(PFWNV1, re.form = ~0, plot = T) # all good
testDispersion(Sim)

PFWNV2 <- lmer(log10(BODY_MASS.x) ~ YEAR + INSECT_T + BROOD + DAYS_TO_HATCH + HATCH_WEIGHTS + (1|NB_YEAR), data=OGRNV)
summary(PFWNV2)
drop1(PFWNV2, test="Chisq");
vif(PFWNV2) # ALL BELOW 4
Sim=simulateResiduals(PFWNV2, re.form = ~0, plot = T) # all good
testDispersion(Sim)

###################
##########                   Pre-fledgling weights Pucon
###################
str(OGRPC)

# Final Model

sapply(D1PCN, function(x) sum(is.na(x) | is.nan(x) | is.infinite(x))) 
D1PCN<- D1PCN[!is.na(D1PCN$RING), ] 

OGRPC <- OGRPC %>%
  left_join(
    D1PCN %>%
      select(RING, HATCH_WEIGHT = BODY_MASS),
    by = "RING"
  )

PFWPC1 <- lmer(log10(BODY_MASS) ~ YEAR + INSECT_T + BROOD + HATCH_WEIGHT + (1|NB_YEAR), data=OGRPC)
summary(PFWPC1)
drop1(PFWPC1, test="Chisq");
vif(PFWPC1) # ALL BELOW 4
Sim=simulateResiduals(PFWPC1, re.form = ~0, plot = T)
testDispersion(Sim)


##### weather differences

t.test(Acc_mill ~ YEAR, data = PCENV) # No
t.test(Av_temp ~ YEAR, data = PCENV) # No
t.test(Min_Temp ~ YEAR, data = PCENV) # No
t.test(Max_Temp ~ YEAR, data = PCENV) # No
t.test(Av_ws_12 ~ YEAR, data = PCENV) # No


M1 <- aov(Acc_mill ~ factor(YEAR), data = NVENV) #No
summary(M1)

M2 <- aov(Av_temp ~ factor(YEAR), data = NVENV) #No
summary(M2)

M3 <- aov(Min_Temp ~ factor(YEAR), data = NVENV) #Yes
summary(M3)
TukeyHSD(M3)

M4 <- aov(Max_Temp ~ factor(YEAR), data = NVENV) #Yes
summary(M4)
TukeyHSD(M4)

M5 <- aov(Av_ws_12~ factor(YEAR), data = NVENV) #No
summary(M5)

TRN <- lm(Av_temp ~ Acc_mill, data=NVENV)
summary(TRN)

plot(Av_temp ~ Acc_mill, data = NVENV,
     col = as.factor(YEAR),
     pch = 16)


NV_1819 <- subset(NVENV, YEAR == 1819)

lm_1819 <- lm(Av_temp ~ Acc_mill, data = NV_1819)
summary(lm_1819)

plot(Av_temp ~ Acc_mill, data = NV_1819,
     main = "1819",
     pch = 16)
abline(lm_1819, lwd = 2)

# Negative relationship between rainfall and temp (higher rainfall = lower temps)

NV_1920 <- subset(NVENV, YEAR == 1920)

lm_1920 <- lm(Av_temp ~ Acc_mill, data = NV_1920)
summary(lm_1920)

plot(Av_temp ~ Acc_mill, data = NV_1920,
     main = "1920",
     pch = 16)
abline(lm_1920, lwd = 2)
# Negative relationship between rainfall and temp (higher rainfall = lower temps)

NV_2023 <- subset(NVENV, YEAR == 2023)

lm_2023 <- lm(Av_temp ~ Acc_mill, data = NV_2023)
summary(lm_2023)

plot(Av_temp ~ Acc_mill, data = NV_2023,
     main = "2023",
     pch = 16)
abline(lm_2023, lwd = 2)
# Negative relationship between rainfall and temp (higher rainfall = lower temps)

PC_1819 <- subset(PCENV, YEAR == 1819)

lm_1819 <- lm(Av_temp ~ Acc_mill, data = PC_1819)
summary(lm_1819)

plot(Av_temp ~ Acc_mill, data = PC_1819,
     main = "1819",
     pch = 16)
abline(lm_1819, lwd = 2)
# Negative relationship between rainfall and temp (higher rainfall = lower temps)

PC_1920 <- subset(PCENV, YEAR == 1920)

lm_1920 <- lm(Av_temp ~ Acc_mill, data = PC_1920)
summary(lm_1920)

plot(Av_temp ~ Acc_mill, data = PC_1920,
     main = "1920",
     pch = 16)
abline(lm_1920, lwd = 2)
```
